# Recovering sedimentary ancient DNA of harmful dinoflagellates off Eastern Tasmania, Australia, over the last 9 000 years

**DOI:** 10.1101/2021.02.18.431790

**Authors:** Linda Armbrecht, Bradley Paine, Christopher J.S. Bolch, Alan Cooper, Andrew McMinn, Craig Woodward, Gustaaf Hallegraeff

## Abstract

Harmful algal blooms (HABs) have significantly impacted the seafood industry along the Tasmanian east coast over the past four decades. To investigate the history of regional HABs, we applied sedimentary ancient DNA analyses (*sed*aDNA) to coastal sediments up to ∼9 000 years old collected inshore and offshore Maria Island, Tasmania. We used metagenomic shotgun sequencing combined with a hybridisation capture array (‘HABbaits1’) to target harmful dinoflagellates of the genera *Alexandrium, Gymnodinium,* and *Noctiluca scintillans*. Bioinformatic analyses were used to verify *sed*aDNA sequences and their presence in older layers, especially for microreticulate cyst forming species including *Gymnodinium catenatum* due to its important role in shellfish toxicity. Our results show that the *Alexandrium* genus (up to 854 and 20 reads per sample inshore and offshore, respectively, based on capture-data) has been present off eastern Tasmania within the last ∼8 307 years. For *G. catenatum* we detected a total of only 9 unambiguously verified reads sporadically between ∼7 638 years ago and the present in the offshore core. We recovered verified *sed*aDNA of the fragile, non-fossilising species *N. scintillans*, along with evidence of increased relative abundance from 2010, consistent with plankton surveys. This study identifies challenges regarding *sed*aDNA sequence validation of some species (in particular, for *G. catenatum*), and provides guidance for the development of tools to monitor past and present HAB species and events, and to improve future HAB event predictions.

**Highlights:**

- Metagenomic *sed*aDNA and hybridisation capture enabled analyses of harmful dinoflagellates off Tasmania
- Sequence validation was used to confirm the presence of *Alexandrium* spp., *Gymnodinium* spp. and *Noctiluca scintillans*
- *Alexandrium* and *Gymnodinium* have been present in Tasmanian waters during the past ∼9 000 years
- *Noctiluca scintillans sed*aDNA derived relative abundance correlates with its recorded increase since 2010

## 1 Introduction

An increasingly important issue is the extent to which harmful algal blooms (HABs) are expanding in distribution, magnitude, and frequency due to climate change, eutrophication, and anthropogenic transport. The negative impacts of HABs on tourism, aquaculture, fisheries, and human health, mean that the appearance of novel HAB phenomena regularly raise the question of whether the species responsible is a recent introduction (e.g., via ballast water; McMinn et al., 1997, Bolch and de Salas 2007), or a previously cryptic endemic species newly stimulated by changing environmental conditions (Hallegraeff et al., 2021) or extreme climate events (Trainer et al., 2019). Few studies have examined the “cryptic species” hypothesis by investigating long-term dynamics over thousands of years (e.g., Thorsen et al., 1995, *Gymnodinium nolleri* misidentified as *G. catenatum* in Scandinavia; Klouch et al., 2016, *Alexandrium minutum* in the Bay of Brest).

The ocean environment off eastern Tasmania is a well-documented climate change hotspot characterised by a strengthening East Australian Current and rapidly increasing ocean temperatures (2.3 °C increase since the 1940s, Ridgway and Hill, 2009). The consequences of this oceanographic change are being detected in coastal marine communities, including changes in plankton and HAB species composition (Thompson et al., 2009, Condie et al., 2019). Particular examples are the HAB dinoflagellates *Gymnodinium catenatum*, *Noctiluca scintillans*, and *Alexandrium* species, which we focus on here.

*G. catenatum* produces Paralytic Shellfish Toxin (PST) and is the only toxic member of a phylogenetically distinct lineage of *Gymnodinium* species (*G. catenatum, G, inusitatum, G. microreticulatum, G. nolleri,* and *G. trapeziforme*) that produce fossilisable resting cysts with distinctive surface reticulation – referred to hereafter as ‘microreticulate species’. Thought to have been introduced to Tasmania in the 1970s by shipping ballast water (McMinn et al. 1997), *G. catenatum* first bloomed in the mid-1980s, caused PST contamination up to 250-fold above acceptable limits, and extensive shellfish farm closures in the Derwent-Huon estuaries from 1986 to 1993 (Hallegraeff et al., 1995). Supporting evidence for a recent introduction includes detection of resting cysts in ships ballast tanks (Hallegraeff and Bolch, 1992), a lack of cysts in dated marine sediment cores prior to the 1970’s (McMinn et al., 1997), and molecular evidence from rRNA gene sequencing that links Australasian populations to source populations from the Seto inland sea in southern Japan (Bolch and de Salas, 2007).

*Noctiluca scintillans* was first documented in Australia from Sydney Harbour (Bennett, 1860), but since the 1990s has increasingly caused highly visible red tides and bioluminescence, resulting in frequent temporary closures of popular Sydney tourist beaches (Hallegraeff et al., 2020). Amongst the suggested causes of *N. scintillans* blooms are eutrophication and coastal upwelling generating more diatom prey (Dela Cruz et al., 2002, 2003). *Noctiluca* was first observed in Tasmanian waters in 1994, and is presumed to have dispersed southward with the East Australian Current, presenting a new threat to the salmonid fish farm industry from 2002 (Hallegraeff et al., 2019). In 2010, *N. scintillans* was detected in the Southern Ocean for the first time, 240 km south of Tasmania, raising concerns of grazing impacts on iconic krill-based food webs (McLeod et al., 2012).

Since 2012, further PST-caused seafood harvest closures of Tasmanian bivalves, abalone, and rock lobsters, along with public health warnings have been generated by winter-spring blooms of the cool-temperate dinoflagellate *Alexandrium catenella.* These events have resulted in PST contamination up to 150 mg saxitoxin (STX) equiv./kg (Condie et al., 2019). The morphologically similar but genetically distinct species *A. australiense* and *A. pacificum* (Bolch and DeSalas, 2007) were previously known from Tasmanian waters, but *A. catenella* (= *A. tamarense* Group 1; =*A. fundyense*; John et al., 2014), had not been previously detected in Australasian waters. Microsatellite DNA analyses of regional populations show that Tasmanian *A. catenella* are divergent from other populations (John et al. 2018), indicating it is either endemic or dispersed naturally to the area millenia ago. Thus, *A. catenella* may be a previously cryptic population recently stimulated by changing environmental conditions, such as increased winter water column stratification (Trainer et al., 2019). At least five cases of non-fatal human paralytic shellfish poisonings (four from *A. catenella* and one from *G. catenatum*) have been formally reported from Tasmania in the past 10 - 20 years (Turnbull et al., 2013; Edwards et al., 2018), but it remains unknown if paralytic shellfish poisonings occurred in this region prior to the arrival of Europeans.

Marine sediment archives containing highly resistant sub-fossil resting cysts are an important resource to examine long-term dynamics of harmful dinoflagellate species and address key questions about their introduction, disappearance, and reappearance in a region. Analysis of sedimentary ancient DNA (*sed*aDNA) from similar samples offers distinct advantages for the analysis of non-cyst forming or less-well preserved cysts that are currently not accounted for in microfossil-based ecosystem reconstructions (Armbrecht, 2020; Bolch, 2022). For example, *Noctiluca* does not produce cysts but has been detected using *sed*aDNA previously (Shaw et al., 2019). However, *sed*aDNA analysis is not straightforward as the DNA is highly fragmented (typically <100 bp), occurs at low concentrations, and any one target species represents only a small fraction of the total DNA present. As a result, it is important to enrich target species *sed*aDNA via methods such as hybridisation capture (Armbrecht, et al., 2021), and to use bioinformatic assessment of DNA damage patterns typical of degraded ancient DNA to authenticate the results (Hübler et al., 2019). Accordingly, an RNA probe array (‘HABbaits1’) was designed previously to bind and retain taxonomic marker genes (18S rRNA, 28S rRNA, Internal transcribed spacer (ITS), ribulose-bisphosphate carboxylase (rbcL), cytochrome c oxidase subunit 1 (COI)) of harmful dinoflagellates (and other plankton groups), including above described *Alexandrium* spp.*, G. catenatum*, and *N. scintillans* (for full details of the HABbaits1 design see Armbrecht et al., 2021).

In this study, we combine metagenomic shotgun sequencing and hybridisation capture using the HABbaits1 array to investigate the presence and palaeo-historical patterns of *Alexandrium, Gymnodinium catenatum,* and *Noctiluca scintillans* from Tasmanian marine sediments over the past ∼9 000 years. We then use this information to support ecological interpretation of regional community change in Tasmanian waters.

## 2 Materials and Methods

### 2.1 Sediment core collection and preparation

An approximately 3 m long sediment core (gravity core, designated ‘GC2S1’) was collected in May 2018 during *RV Investigator* voyage IN2018_T02 in 104 m water depth close to the continental shelf edge, east of Maria Island, Tasmania (Site 1; 148.240 °E; 42.845 °S) (Fig. 1). Since gravity cores can disturb surface sediments, and the latter can be lost due to the horizontal tipping after collection, a shorter (12 cm) parallel core was obtained at the same site using a KC Denmark Multi-Corer (designated ‘MCS1-T6’, where ‘T6’ refers to ‘Tube 6’ of the multi-corer). A 35 cm long multi-core was also obtained from within Mercury Passage, west of Maria Island (Site 3; 42.550 °S, 148.014 °E) in a water depth of 68 m (designated ‘MCS3-T2’). All cores were immediately capped, sealed, labelled and transported in their original PVC coring tubes to the Australian Nuclear Science and Technology Organisation (ANSTO), Lucas Heights, Australia, where they were kept at 4°C. The cores GC2S1, MCS1-T6, and MCS3-T2 were opened, split, scanned (using a multi-function core scanning instrument (ITRAX) with X-ray fluorescence (XRF), radiographic X-ray, optical imaging and magnetic susceptibility measurements), and subsampled for *sed*aDNA analyses in October 2018 (i.e., as fast as possible considering the logistics involved). To minimise contamination during core splitting and sampling, working benches and cutting knives were cleaned with 3% bleach and 70% ethanol, gloves changed immediately when contaminated with sediment, and appropriate PPE was worn at all times (gloves, facemask, hairnet, disposable coverall). Sampling of GC2S1 was conducted by first removing the outer ∼1 cm of the working core-half and then taking subsamples by pressing sterile 15 mL centrifuge tubes ∼3 cm into the sediment at 5 cm intervals (working from bottom to the top of the core at each step). Sampling of MCS1-T6 was conducted as for GC2S1 except at finer intervals of 2 cm. Mercury Passage multicore (MCS3-T2) sampling was conducted at 2 cm depth intervals in the top 8 cm and at 5 cm intervals below. All *sed*aDNA samples were immediately stored at -20 °C, respectively. Hereafter, we refer to sediment depths as 0, 2 cm, *etc.*, however it should be noted that due to the diameter of the sample collection centrifuge tubes (∼1.5 cm diameter), this refers to the minimum depth of the sample interval which was actually 0 - 1.5 cm, 2 – 3.5 cm, etc., respectively.

**Figure 1:**
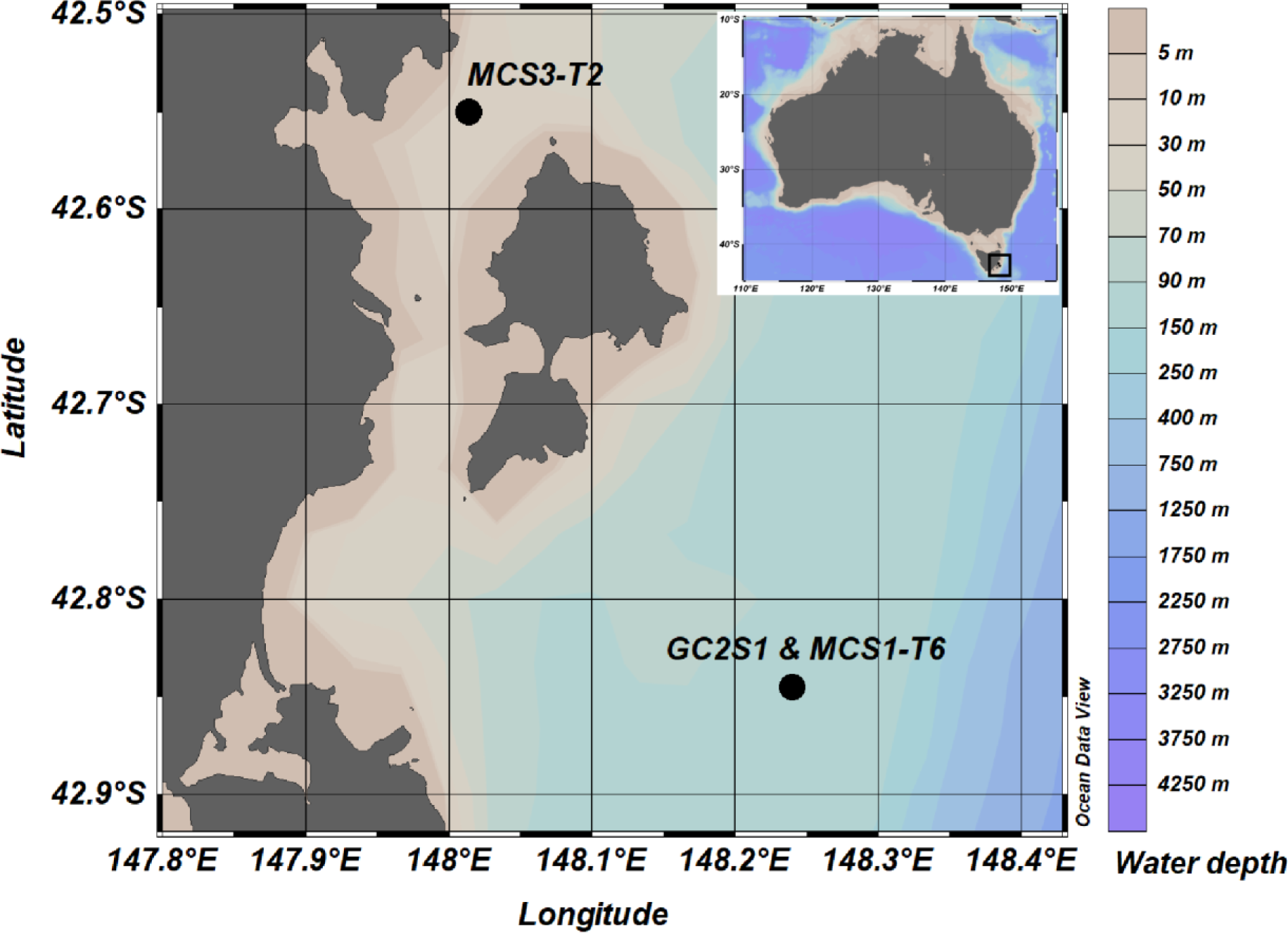
Sediment coring sites near Maria Island, Tasmania, Australia. Overview of coring locations offshore (Gravity Core Site 1, GC2S1, and Multi-Core Site 1 Tube 6, MCS1-T6) and inshore Maria Island in the Mercury Passage (Multi-Core Site 3 Tube 2, MCS3-T2). GC2S1 and MCS3-T6 were collected adjacent to each other and are depicted as one dot. Map created in ODV (Schlitzer, R., Ocean Data View, https://odv.awi.de, 2018).

### 2.2 Sediment dating

ITRAX scanning, XRF, radiographic X-ray, optical imaging and magnetic susceptibility measurements confirmed excellent undisturbed preservation of the cores. An age profile was generated for MCS3-T2 and MCS1-T6 based on ^210^Pb measurements (8 and 6 dates, respectively), while GC2S1 was based on both ^210^Pb (7 dates) and ^14^C (3 dates). A Bayesian age-depth model was constructed for each site based on these ^210^Pb and ^14^C measurements using rbacon (Blaauw et al., 2019) on the R platform (R Core Team, 2013) with the SHCal20 curve for radiocarbon age calibration (Hogg et al., 2020). Details on the construction of the age-depth model are provided within the Supplementary Material (Supplementary Material Note 1, Supplementary Material Fig. 1, Supplementary Material Table 1).

### 2.3 Sedimentary ancient (sedaDNA) extractions

Extractions of *sed*aDNA took place within ultraclean ancient (GC2S1) and forensic (MCS1-T6, MCS3-T2) facilities (ACAD, The University of Adelaide) following recommended ancient DNA decontamination standards (Willerslev and Cooper, 2005). The extraction technique followed the ‘combined’ protocol developed for marine eukaryote *sed*aDNA (Armbrecht et al., 2020). This method combines a gentle EDTA incubation step to isolate fragile eukaryote DNA (Slon et al., 2017) with bead-beating to extract intracellular DNA from robust spores and cysts (Shaw et al., 2019). Smaller DNA fragments (≥27 base pairs, bp) characteristic of ancient DNA are targeted using in-solution silica binding (Brotherton et al., 2013), along with magnetic beads to size-select DNA fragments under 500 bp (Armbrecht et al., 2020). DNA extracts were prepared from 42 sediment samples (GC2S1, MCS1-T6, MCS3-T2) and 7 extraction blank controls (see also Armbrecht et al., 2021).

### 2.4 Shotgun sequencing library preparations

Metagenomic shotgun libraries were prepared from 42 sediment samples (GC2S1, MCS1-T6, MCS3-T2) and 7 extraction blank controls using established techniques (Armbrecht et al., 2020, 2021). Sequencing was undertaken using an Illumina NextSeq sequencing platform (2 x 75 bp cycle) at the Australian Cancer Research Foundation Cancer Genomics Facility & Centre for Cancer Biology (Adelaide, Australia) and the Garvan Institute of Medical Research, KCCG Sequencing Laboratory Kinghorn Centre for Clinical Genomics (Darlinghurst, Australia).

### 2.5 Hybridization capture and sequencing library preparations

In order to maximise the yield of *sed*aDNA from our target dinoflagellate species a hybridization-capture technique was developed and applied. An RNA array or ‘bait set’ termed ‘HABbaits1’ was developed in collaboration with Arbor Biosciences, USA, to target the harmful dinoflagellates *Alexandrium* groups I – IV, *Gymnodinium catenatum* and *Noctiluca scintillans*. Details of the design, protocol optimisation, and application of HABbaits1 to marine *sed*aDNA have been published previously (Armbrecht et al., 2021). A final pool of a selection of 30 multiplexed sequencing libraries (GC2S1 and MCS3-T2) prepared from HABbaits1 was sequenced using Illumina HiSeq XTen (2 x 150 bp cycle) at the KCCG, Darlinghurst, Australia.

### 2.6 Bioinformatics

Bioinformatic processing of sequence data followed published protocols (Armbrecht et al. 2020), using software and analytical parameters from Armbrecht et al. (2021). After filtering to remove low-complexity and duplicate reads, each dataset was processed without the standardisation step (*i.e.,* without rarefying) to retain the maximum number of reads, which is crucial for ancient DNA damage analysis (see below). However, to assess our data semi-quantitatively, we also standardised the filtered shotgun data by subsampling (*i.e.,* rarefying) to the lowest number of reads detected in a sample, *i.e.,* 2.2 million (Mio) (using seqtk version 1.2), and continued to process this data in parallel to the non-rarefied data. Subsampling reduced the number of reads significantly therefore this data is provided in the Supplementary Material only.

After quality control (FastQC v.0.11.4, MultiQC v1.8), the NCBI Nucleotide database (ftp://ftp.ncbi.nlm.nih.gov/blast/db/FASTA/nt.gz, downloaded November 2019) was used as the reference database to build a MALT index (Step 3) and sequences aligned using MALT (version 0.4.0; semiglobal alignment) (Herbig et al., 2016). All resulting .blastn files were converted to .rma6 files using the Blast2RMA tool in MEGAN (version 6_18_9, Huson et al., 2016). Subtractive filtering (*i.e.,* subtracting reads for species identified in EBCs from samples) was conducted separately for the shotgun and HABbaits1 data (see Armbrecht et al., 2021 Supplementary Material, for a comprehensive list of eukaryote contaminants), however, no Dinophyceae taxa were detected in EBCs. For both datasets, read counts of all Dinophyceae nodes (from MEGAN6 v. 18.10) were exported for downstream analyses.

To test *sed*aDNA damage, the ‘MALTExtract’ and ‘Postprocessing’ tools of the HOPS v0.33-2 pipeline (Hübler et al., 2020) were run for our three target dinoflagellates using the same configurations as used in Armbrecht et al. (2021), eg. taxalist *‘b’*, including *Alexandrium* spp., *Gymnodinium* spp., and *N. scintillans* on shotgun and HABbaits1 *sed*aDNA data. The MALTExtract output, *i.e.,* reads categorised as ancient (showing damage) or default (passing stringent filtering criteria but not showing damage) for the three target taxa was exported and the proportion of *sed*aDNA damage per taxon was determined. Also, *sed*aDNA damage profiles were generated for these three taxa using MALTExtract Interactive Plotting Application (MEx-IPA, https://github.com/jfy133/MEx-IPA), however, due to low read numbers these graphics were not informative and excluded from downstream analyses.

To test the reliability of short sequence *sed*aDNA assignments to the three targets groups, we created three dummy sample datasets, each containing sequences of one of the three dinoflagellate groups downloaded from NCBI (see Supplementary Material Note 3). Each sequence was included with its complete length (Supplementary Material Table 2) and split into 56 bp fragments (corresponding to the average sequence length of our filtered shotgun data, see Results). If the last fragment was <25 bp it was excluded from the dummy sample (mimicking a minimum cut-off of 25 bp during sample data processing). Dummy samples were converted from .fasta to .fastq files (see Supplementary Material Note 3) and processed via the same analytical pipeline used for sample sequencing.

The initial assessment of s*ed*aDNA assigned to *Gymnodinium catenatum* indicated a significant proportion of sequences mapped to regions of the LSU-rRNA gene where there is limited database coverage of gymnodinoid dinoflagellates (see Supplementary Material Note 4, Supplementary material Figs. 4,5). Due to the potential for reference bias, we undertook additional sequencing of the D3 to D10 region (∼1 850 bp) of the LSU-rRNA of related microreticulate species available to us in culture, *G. microreticulatum* (strain CAWD191) and *G. nolleri* (strain K-0626) (Supplementary Material Note 4) and carried out more detailed validation of these sequences Firstly, gymnodinoid-assigned reads were mapped to *G. catenatum* reference sequence (DQ785882) including comparative alignment with related *Gymnodinium* species (default alignment and assembly; Geneious Prime 2021). Direct match/mismatches to species in the alignment were used to determine the nearest-neighbour taxon and base-pair mismatches from the reference sequence for each read (Supplementary Material Fig. 5). Secondly, all *Gymnodinium* reads assigned to *G. catenatum* after the initial NCBI run were then re-run against an in-house *Gymnodinium*-focused reference sequence database with addition of newly generated *G. microreticulatum* and *G. nolleri* LSU-rRNA D3-D10 sequences (see Supplementary Material Note 5).

## 3 Results

### 3.1 Age model

Our age model revealed that the inshore core MCS3-T2 dates to 77 years before 1950 at 34.5 centimetres below seafloor (cmbsf), *i.e.,* the year 1873 (mean value, ∼144 years before sample collection in 2017). The offshore gravity core GC2S1 dates to 8 878 years before 1950 at 268 cmbsf (∼8 945 years before sample collection in 2017). Comparing the ^210^Pb activity profiles from both GC2S1 and the adjacent multicore MCS1-T6, we estimated that ∼3.5 cm were missing from the top of GC2S1, representing the last ca. 30 years (Supplementary Material Fig. 1).

### 3.2 Validation of Alexandrium, Gymnodinium., and Noctiluca scintillans short read species assignments

Dummy dataset analyses revealed short reads of *Alexandrium* spp. could be reliably assigned to genus level, thus we interpret this target taxon exclusively at genus level (see Supplementary Material Note 3, Supplementary Material Table 2, Supplementary Material Fig. 3). For *N. scintillans*, the majority of dummy sequences were correctly assigned to this species, and correctly back-mapped to the NCBI sequence used to create our dummy sample (GQ380592.1). Some reads were assigned to a different *N. scintillans* 18S rRNA reference sequence available on NCBI (AF022200.1), but species-level assignment remained correct therefore we consider identification of *N. scintillans* from short *sed*aDNA fragments to be robust and reliable. For *Gymnodinium catenatum*, most dummy reads were correctly assigned but few reads were erroneously assigned to the apicomplexan genus *Eimeria*, the Streptophytina, the Membracoidea, other dinoflagellates such as *Durinskia baltica, Symbiodinium* sp. AW-2009, or the related microreticulate species *G. microreticulatum*, suggesting *Gymnodinium* species assignment of short reads (56 bp) of 18S rRNA should be interpreted with caution. In most cases assignment uncertainty could be resolved by further inspection and verification. Similarly, rare mis-assignment of short 18S rRNA reads from *Alexandrium* were made to Apicomplexa and Apocrita, and for *N. scintillans* to *Symbiodinium* and Apocrita.

Reference alignment mapping of HABbaits1 reads assigned to *Gymnodinium* from core GC2S1 (352 reads, see Results), showed the majority (99%) mapped to the rRNA genes (Supplementary Material Note 4, Supplementary Material Fig. 5). Of these, 65% were within 3 bp of exact match for the *G. catenatum* reference sequence (Supplementary Material Table 3) and 97% could be unambiguously assigned to the microreticulate-group of *Gymnodinium* species that includes *G. catenatum*. More than half (57%) of the reads were closest to the *G. catenatum* reference sequence, 36% unambiguously identified as *G. catenatum*, and 12.5% as *G. microreticulatum* (Supplementary Material Fig. 5). Twenty-one percent mapped to rRNA regions where *G. catenatum* could not be resolved from its closest relative *G. nolleri*; 30% to regions with insufficient sequence variation to distinguish microreticulate-group species (Supplementary Material Figure 5).

Re-assignment of the 352 gymnodinoid reads against our supplemented in-house database supported these observations and resulted in more conservative species-level assignment. Of 216 reads assigned, 76% could only be assigned at genus level, 28% to *G. nolleri*, 6% to *G. microreticulatum*, and 4% to *G. catenatum* (Supplementary Material Note 5, Supplementary Material Table 4).

### 3.3 Representation of Dinophyceae in shotgun data

From the 42 shotgun samples we retrieved a total of 824 503 filtered sequences across the three domains (Bacteria, Archaea and Eukaryotes), 149 892 of which were assigned to Eukaryota (18%). A total of 529 reads were assigned to Dinophyceae (*i.e.,* Dinophyceae represented 0.06% of the domains, and 0.35% of all eukaryotes). Harmful dinoflagellate taxa sequences were detected in low abundance (19 *Alexandrium* spp., 13 *Gymnodinium* spp., and no *Noctiluca* sequences; Fig. 2). *Alexandrium* spp. were primarily detected in the upper 15 cmbsf at MCS3-T2 (last ∼39 years), and few reads were assigned to this genus between 55 and 189 cmbsf at GC2S1 (∼2 600 and 7 200 years ago, respectively). Based on our *Alexandrium* dummy run (Section 3.2) we used a conservative approach, at genus level (Fig. 2). Three sequences were assigned to microreticulate *Gymnodinium* group in GC2S1 at 95 cmbsf (∼4,400 years ago), and 219 cmbsf (∼7,800 years ago) (Fig. 2C). While microreticulate gymnodinoids were not detected in the younger sediments of GC2S1, they were present in the surface sediments of the shorter MCS1-T6 core (Fig. 2B), demonstrating their presence in more recently deposited offshore sediments.

**Figure 2:**
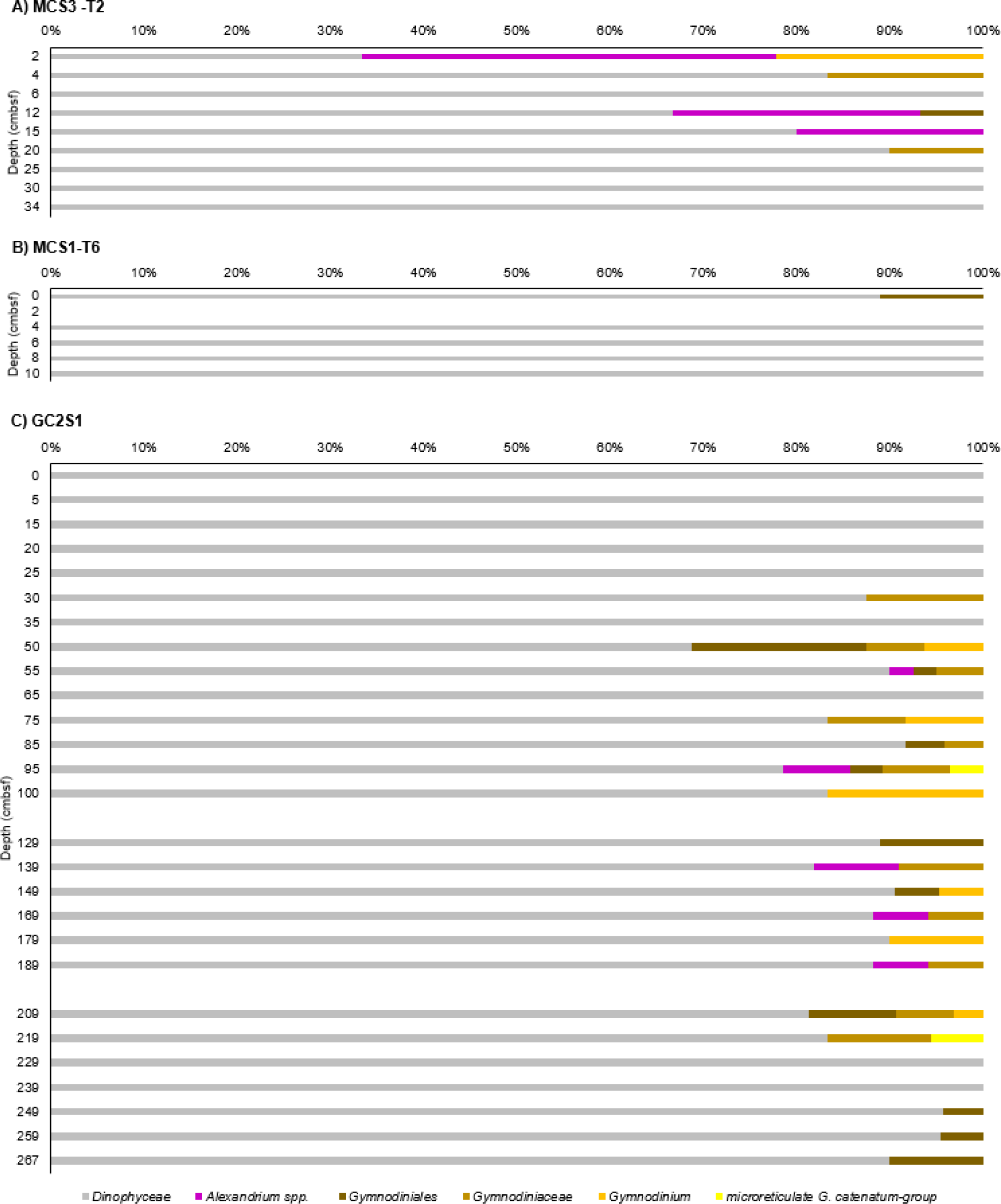
Relative Dinophyceae abundance (shotgun). Dinophyceae detected in non-rarefied shotgun data at Maria Island coring sites A) MCS3-T2, B) MCS1-T6, and C) GC2S1. All non-target taxa are grouped as Dinophyceae (grey), while *Alexandrium* and Gymnodiniales and its subgroups are shown separately (in colour). Identity of *G. catenatum* should be interpreted as representing all microreticulate *Gymnodinium* species, and the genus *Alexandrium* (see Section 3.2). *N. scintillans* was not detected in the shotgun data. Total read count (all sites): 529.

### 3.4 Representation of Dinophyceae using HABbaits1

Hybridisation capture of our 30 selected *sed*aDNA extracts with the HABbaits1 array resulted in 27 successfully enriched samples from GCS1 and MCS3-T2 with a total of 872 774 reads across the Bacteria, Archaea and Eukaryote domains, of which 613 853 were assigned to Eukaryota (70%). A total of 32 075 of the eukaryote reads were assigned to Dinophyceae (*i.e.,* 3.68% of all domains, and 5.23% of all eukaryotes), representing a 61- and 15-fold increase in Dinophyceae reads relative to shotgun data, demonstrating the efficacy of target enrichment achieved with HABbaits1.

The genus *Alexandrium* were most abundant at MCS3-T2, with the maximum number of reads (854 sequences) found inshore at MCS3-T2 12 - 13.5 cmbsf (∼190 years ago) (Fig. 3A). At GC2S1, *Alexandrium* spp. were detected sporadically at different depths at low abundance (<20 reads per sample) (Fig. 3B).

**Figure 3:**
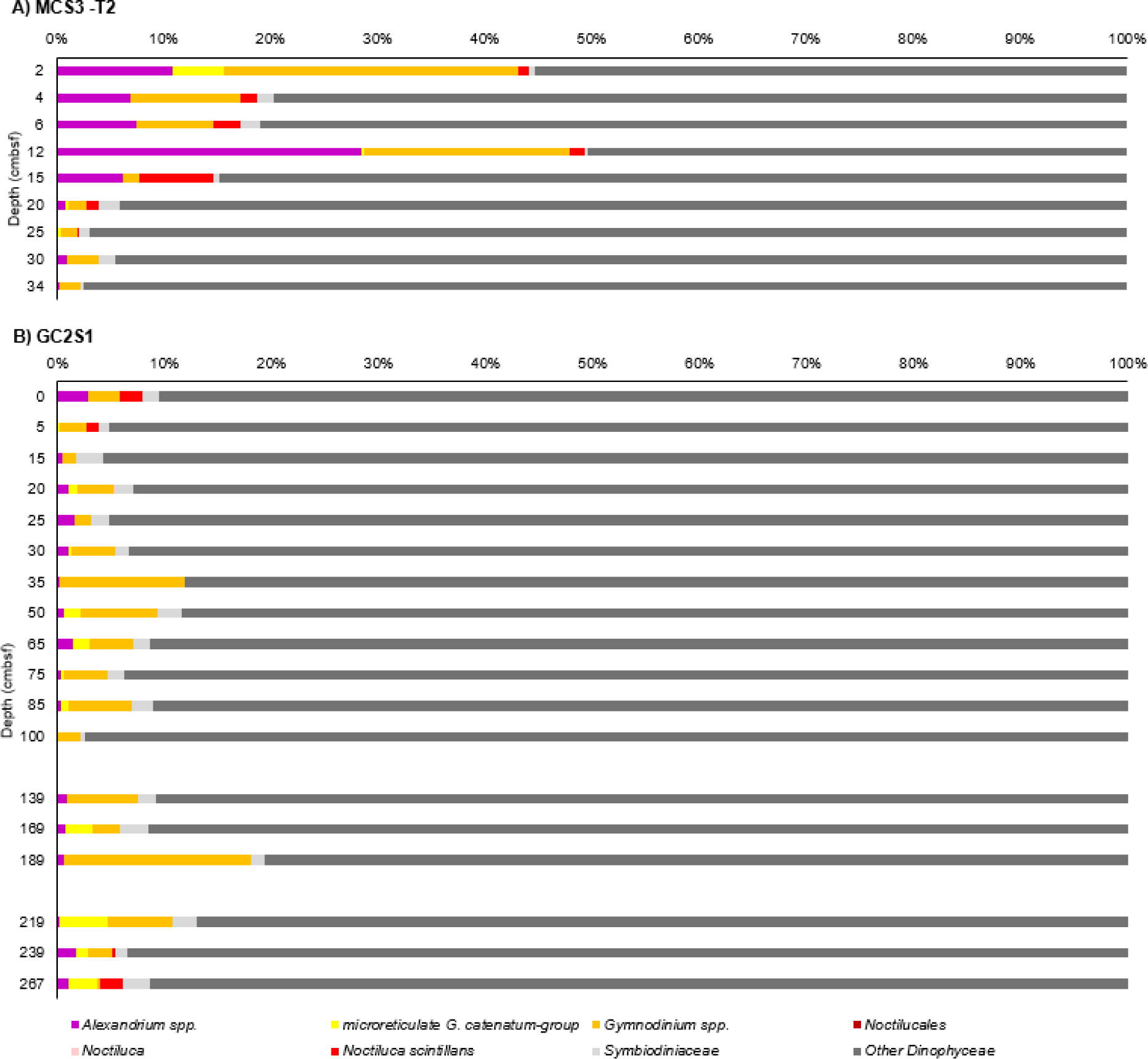
Relative Dinophyceae abundance (HABbaits1). Dinophyceae detected in non-rarefied HABbaits1 data at Maria Island coring sites A) MCS3-T2 and B) GC2S1, with target taxa highlighted. Highlighted are *Alexandrium* spp. (pink), the microreticulate gymnodiniods (yellow) and broader genus *Gymnodinium* spp. (orange), and *N. scintillans* (red) within the broader group Noctilucales/*Noctiluca* (total of 1 and 2 reads only, respectively). All other Dinophyceae are summarised (dark grey), with *Symbiodiniaceae* shown separately (light grey) due to potential misassignments with *Gymnodinium* or *Noctiluca* (Section 3.2). Total number of reads (both sites): 32 075.

Several reads were assigned to the microreticulate *G. catenatum*-group at both MCS3-T2 and GC2S1 (352 reads in total). A relatively high abundance (85 reads) of this group was identified inshore in MCS3-T2 2 - 3.5 cm (∼2 years ago, i.e., 2015), while at GC2S1, the microreticulate *G. catenatum*-group appeared sporadically offshore in low abundance (≤22 reads) in the upper section of GC2S1 (above 75 – 76.5 cmbsf; ∼3,500 years ago), while being slightly more abundant (≤123 reads) in the lower core section (below 169 – 170.5 cmbsf, ∼6,500 years ago; Fig. 3B). Re-assignment of the 352 reads initially identified as *G. catenatum* to the new *Gymnodinium* database provided 216 reads assigned to *Gymnodinium* spp. (all others remained unassigned). Most reads were assigned on genus level (165 reads), and 28, 14 and 9 reads were assigned to *G. nolleri, G. microreticulatum* and *G. catenatum,* respectively (Fig. 4). *G. nolleri* was identified sporadically through GCS1, while *G. microreticulatum* and *G. catenatum* were found primarily in surface sediments at MCS3 and/or GC2S1 as well as sporadically in deeper samples at GC2S1 (e.g., 209 cmbsf, ∼7,638 years ago) (Fig. 4).

**Figure 4:**
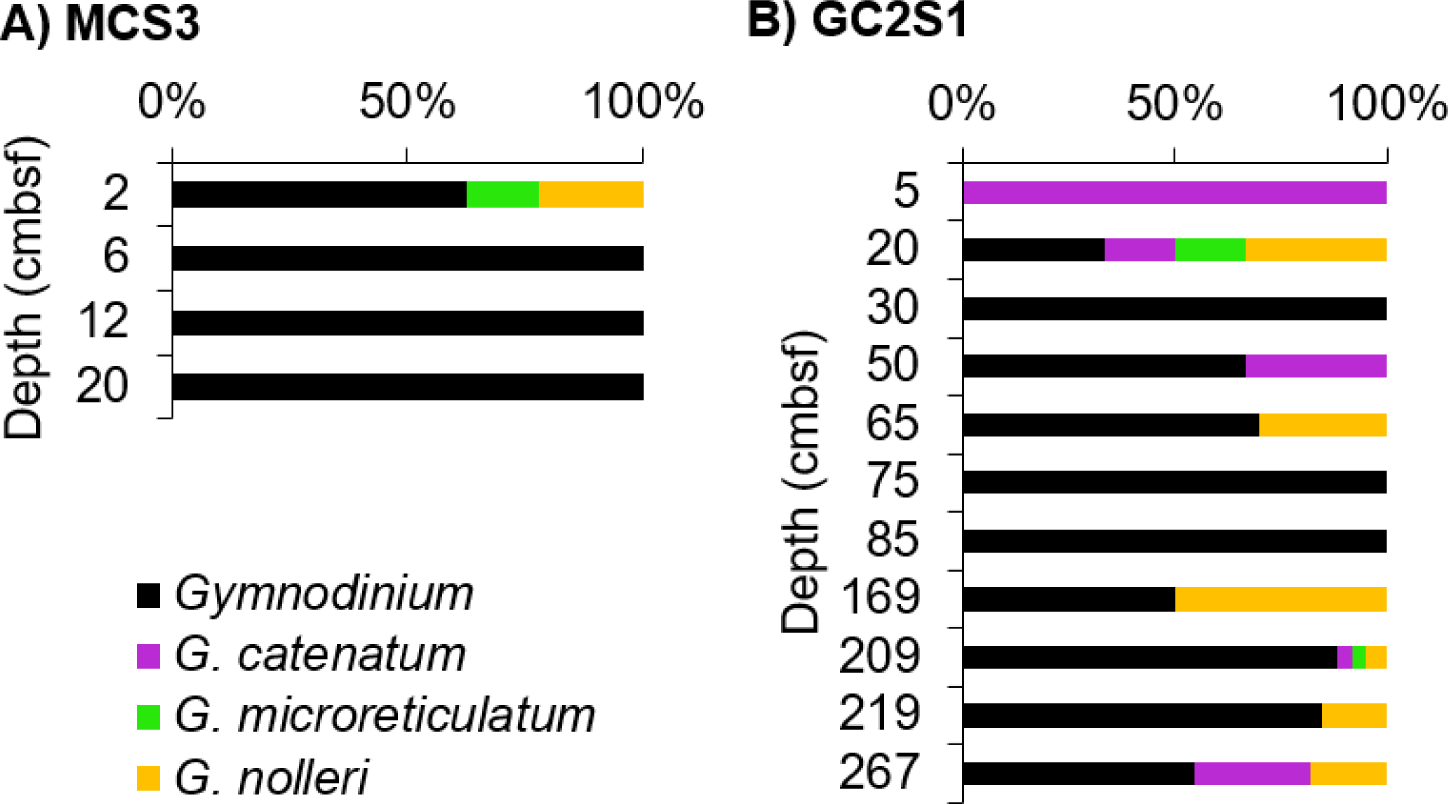
Microreticulate *Gymnodinium* species. Re-assignment of 352 *Gymnodinium* spp. at Site MCS3 (A) and GC2S1 (B) using an in-house *Gymnodinium* spp. database supplemented with additional microreticulate rRNA gene sequences. A total of 216 reads were assigned to genus and/or species level. The deepest sample at GC2S1 (267 cmbsf) should be interpreted with caution due to potential seawater contamination during core retrieval.

Hybridisation capture also allowed the identification of *N. scintillans*, primarily in the upper section of MCS3-T2 (above 15 cmbsf), reaching maximum relative abundance (based on 59 sequences) at 6 - 7.5 and 12 - 13.5 cmbsf (∼7 and 15 years ago, *i.e.,* around 2010 and 2002) (Fig. 3A). *N. scintillans* was also detected in the two top samples at GC2S1 (≤8 reads each, Fig. 3B).

All three target dinoflagellate taxa were detected in low relative abundance in the deepest sample (Fig. 3B), but we cannot rule out the possibility of seawater contamination of the bottom core sample during core retrieval. Additionally, dummy sample validation indicates the possibility that shorter *Gymnodinium* and *Noctiluca* sequences may be misassigned to *Symbiodinium* (see Section 2.3), increasing relative abundance of these taxa by ∼1% (Fig. 3).

### 3.5 sedaDNA damage analysis and authentication

Application of the HOPS DNA damage approach to the HABbaits1 data at site MCS3-T2 identified 30, 4, and 1 reads with ancient characteristics (*i.e.,* showed clear signs of DNA damage) for *Alexandrium, G. catenatum*, and *N. scintillans*, respectively. At site GC2S1, and 22, 95, and 11 ‘ancient’ reads were identified for *Alexandrium*, *G. catenatum*, and *N. scintillans*, respectively (Table 1). A relatively high number of reads passed the default filtering criteria in HOPS (Table 1) - 2 and 15% of the *Alexandrium* reads were classified as ancient in MCS3-T2 and GC2S1, respectively (Fig. 5). For *G. catenatum*-group, 4 and 27% of reads were classified as ancient in MCS3-T2 and GC2S1, respectively, and for *N. scintillans*, 0.5 and 18% were classified as ancient (Table 1, Fig. 5). As such, reads assigned to the three target dinoflagellates showed much higher *sed*aDNA damage at the offshore site GC2S1 relative to the inshore site MCS3-T2.

**Figure 5:**
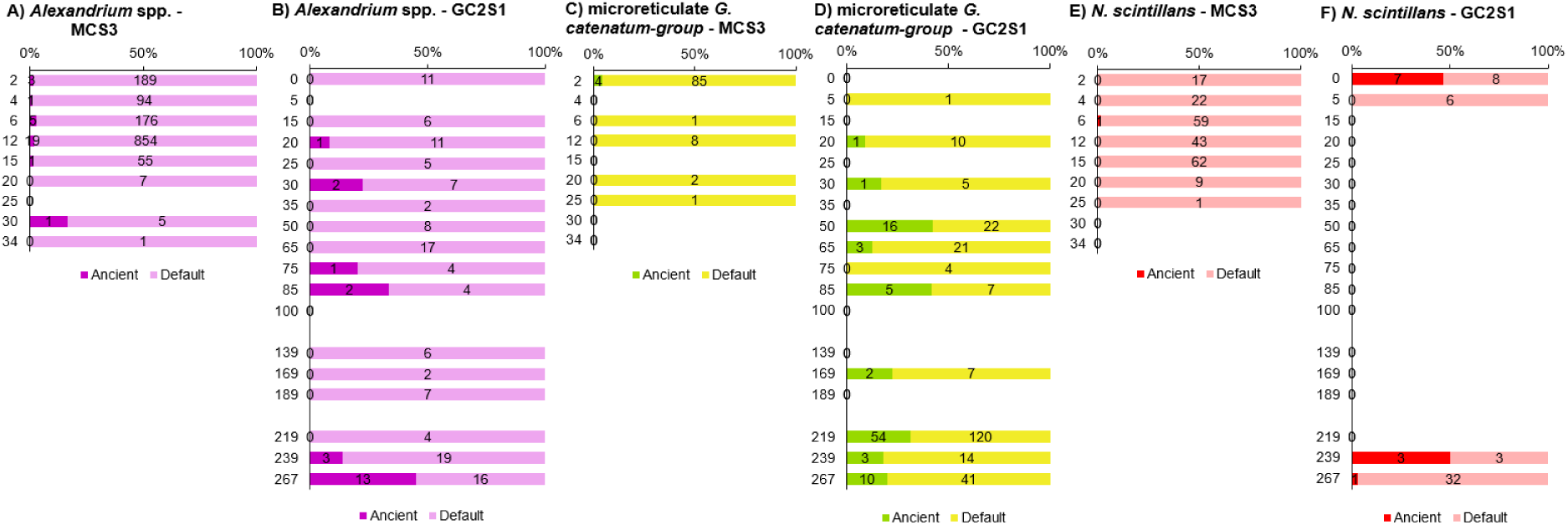
Ancient and default reads identified for three target HAB taxa in MCS3-T2 and GC2S1. Shown are the proportion of *sed*aDNA reads classified by HOPS analysis as ancient and default (% DNA damage) per sample based on the HABbaits1 capture sequences. The number of reads underlying these proportions are indicated within the bars.

**Table 1:**
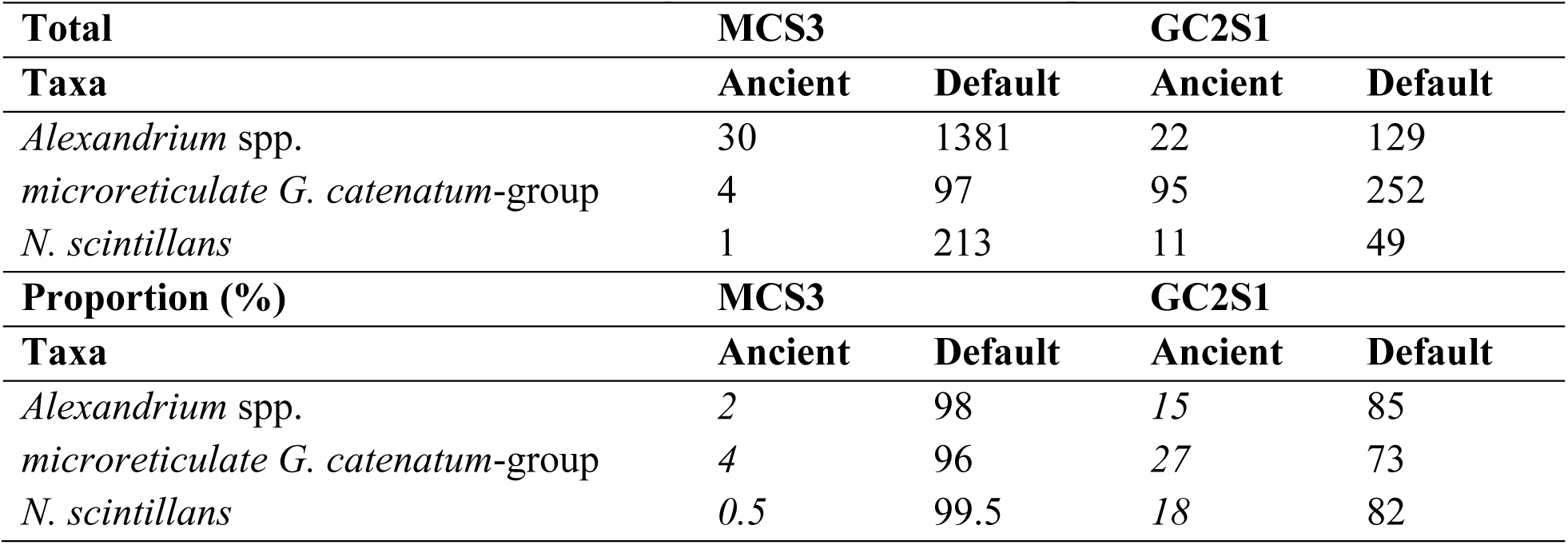
*sed*aDNA damage of reads assigned to *Alexandrium* genus, *G. catenatum*-group and *N. scintillans*. The total number and proportion of reads classified into ancient and default via HOPS *sed*aDNA damage analysis per site (based on HABbaits1). The proportion of ancient reads is a measure of ‘% *sed*aDNA damage’ for each of the three species (in italics).

## 4 Discussion

Our analysis of dinoflagellate *sed*aDNA shows that ancient DNA from the genera *Alexandrium* and *Gymnodinium* is preserved in marine sediment for thousands of years. In contrast, *N. scintillans* DNA was detected primarily in recent sediments. The up to 60-fold increase in dinoflagellate sequences achieved using hybridisation capture enrichment with the HABbaits1 array demonstrates the value of this approach for HAB focused *sed*aDNA studies. Additional optimisation, such as adjusting the hybridisation protocol (hybridisation temperature and time, Horn et al., 2012; MyBaits, 2018), and increased sequencing depth, will make it possible to maximise the ancient genetic signal of individual marine species over millennial timescales.

### 4.1 Considerations for interpretation of Alexandrium, Gymnodinium and Noctiluca sedaDNA detection

Due to the short length of ancient DNA reads (mean ∼56 bp; shotgun data post-filtering), we expected a degree of uncertainty in their assignment to reference sequences, at a species level. Our dummy data processing runs containing fragments of taxonomic marker genes of the three target dinoflagellates *Alexandrium* spp., *Gymnodinium* spp., and *N. scintillans* provided a useful means to assess the level of species assignment uncertainty on a taxon-by-taxon basis. For *Alexandrium,* the dummy sample analyses indicated our short sequences could not be confidently assigned to species, resulting in our use of a conservative genus-level interpretation. In Tasmanian waters, this genus includes *A. affine, A. australiense, A. catenella, A. margalefi, A. ostenfeldii, A. pacificum* and *A. pseudogonyaulax* (Hallegraeff et al. 1991, Bolch & de Salas 2007, Ruvindy et al. 2018). Resolving this complex at species or even genotype-level could provide valuable information on the history and distribution of toxic members of this genus. However, given the low confidence in assigning the *Alexandrium* dummy sequences below genus level, this is beyond the scope of this study and would require optimisation of RNA bait-design, hybridisation capture (*e.g.*, increased input material), and/or gene assembly to generate longer DNA fragments, in order to achieve more reliable taxonomic assignments.

It was possible to make more confident species level assignments for *N. scintillans* due to the uniquely monospecific nature of this genus (Gómez et al., 2010). The few mis-assignments noted were 18S rRNA sequences of our target dinoflagellates (Supplementary Material Note 3, Supplementary Material Fig. 3), reinforcing our conclusions that this marker gene is too conserved to confidently assign very short dinoflagellate reads. While the HABbaits1 design included multiple marker genes for harmful dinoflagellates occurring off Tasmania, future RNA bait designs focused on the LSU-rRNA and rRNA-ITS are likely to be more effective and lead to higher confidence in species-level assignment of *sed*aDNA from this region (and other locations; Bolch, 2022).

The target gene regions included in the HABbaits1 array were sufficiently variable to distinguish gymnodinoid dinoflagellates, including the microreticulate group of species that includes *G. catenatum*. While exact sequence matches to *G. catenatum* reference sequence DQ785882 were rare, >60% were within 3 bp of this reference, and in regions with sufficient variation to distinguish between microreticulate-group species, including the most closely related *G. nolleri* and *G. microreticulatum* (Supplementary Material Fig. 5, Supplementary Material Table 3). The reference sequence DQ785882 was derived from a Korean *G. catenatum* isolate, therefore, the observed differences may partly be attributed to global variations in rRNA sequence of *G. catenatum* (Bolch and De Salas, 2007; Band-Schmidt et al. 2008). An alternative is that processed (de-duplicated) *sed*aDNA data effectively represents a single fragment from the original genome of the target species. Dinoflagellate genomes possess many thousands to millions of rRNA gene loci (Ruvindy et al., 2023) considerable intra-genomic sequence heterogeneity (Gribble and Anderson, 2006; Miranda et al., 2012), multiple sequence classes and the presence of pseudogenes (Scholin and Anderson, 1994). Numerous low frequency sequence variants have also been detected in cultures established from single cells (Miranda et al. 2012). As a result, the single-based variation we noted among *sed*aDNA reads may result from the *sed*aDNA pipeline faithfully retaining different single nucleotide mutations at different rRNA loci from the original genomes.

Both our dummy data and manual fragment mapping indicated an assignment to the microreticulate group of gymnodinoid species at high confidence, but the resolution of the five species is partly dependent on both fragment length and gene region/location of each *sed*aDNA read. The net effect is that only a proportion (up to 33%) of microreticulate species reads could be unambiguously assigned to an individual species, but that those that could be assigned to species-level using the *Gymnodinium*-only database are robust. Based on our validation studies (Supplementary Material Notes 4,5), HABbaits1 can be feasibly applied to gymnodinoids more generally, and other dinoflagellate genera with similar levels of rRNA-ITS and LSU-rRNA sequence variation between species. The mapping to reference sequences also highlighted the potential for assignment artefacts arising from limited database coverage. Robust species assignments of microreticulate gymnodinoids, including toxic *G. catenatum*, was only possible after addition of the new D3-D10 LSU-rRNA reference sequences. The improved target gene coverage in the database allowed detection of the non-toxic related species *G. microreticulatum*, which is uncommon but widely distributed in both Australian and Tasmanian waters (Bolch and Hallegraeff, 1990; Bolch and Reynolds, 2002), and sequences assigned to *G. nolleri* which had not been previously identified from the Australasian region by microscopy. While *G. nolleri* appears confined primarily to coastal European waters, it is conceivable this species is distributed very widely but is rare/cryptic in Tasmanian waters. It may also have been potentially more abundant in the past, as shown in the Baltic Sea where increased abundance of *G. nolleri* resting cysts coincided with the Medieval Warm Period, affecting the north Atlantic region from circa 800-1250 AD (Thorsen et. al., 1995).

The *sed*aDNA approach used here allowed the authenticity of the ancient sequences recovered for the three target taxa to be assessed. While % *sed*aDNA damage was low (≤4%) at the inshore site MCS3-T2, this is consistent with the earlier studies showing that eukaryote *sed*aDNA damage is very low in the upper ∼35 cm of sediment at this site (Armbrecht et al., 2021). At GC2S1 (longer offshore core), the increased *sed*aDNA damage observed (up to 27% for *G. catenatum*) indicate that the recovered sequences from *Alexandrium*, the microreticulate gymnodinoids and *N. scintillans* are consistent with authentic *seda*DNA, except for the samples from the bottom of the core, which were potentially contaminated by seawater during core retrieval.

### 4.2 Alexandrium

The distribution of *Alexandrium sed*aDNA reads throughout the offshore core, including the deepest samples, indicates that *Alexandrium* has been present in the area during the past ∼9 000 years. The HABbaits1 data also show that *Alexandrium* has recently increased in relative abundance in inshore waters (from ∼15 years ago; circa 2003), but has remained in low relative abundance offshore (<3%).

Compared to microreticulate gymnodinoid species that are typically rare in coastal sediments (<2% of cysts; Bolch and Hallegraeff, 1990), *Alexandrium* species can be prolific producers of resting cysts (approximately 40% of vegetative cells; Anderson et al., 2014). Live resting cysts are resistant to chemical and biological attack, protecting DNA from degradation during transport to the bottom sediment. Some cysts can remain dormant but viable in anoxic sediments for >100 years (Feifel et al., 2012), indicating functional and undegraded DNA survival for centuries. The patterns of abundance observed in *sed*aDNA mirror trends observed in microscopy-based counts of *Alexandrium* resting cysts from the same cores (Paine et al., in review). The highest *Alexandrium* cyst concentrations were observed in the top 35 cmbsf at MCS3-T2 (inshore), while low concentrations of *Alexandrium*-like cysts were observed throughout the GC2S1 core (offshore). Very few *Alexandrium* sequences were detected offshore, but the trend towards increased *sed*aDNA damage with depth (especially at GC2S1) supports the authenticity of the ancient reads.

Unfortunately, our HABbaits1 enriched *seda*DNA was unable to discriminate between *Alexandrium* species (see section 4.1). However, the relative abundance patterns support the view that toxic *Alexandrium* species (*A. catenella* and *A. pacificum*) are endemic in the Tasmanian region, but are cryptic, low abundance species not previously distinguished from morphologically similar and generally non-toxic *A. australiense*. Their recent increased relative abundance in inshore cores also support the view that emergence of blooms from 2012 onwards have been stimulated by climate-induced changes in environmental conditions (Condie et al., 2019).

### 4.3 Gymnodinium catenatum and microreticulate species

The HABbaits1 array detected microreticulate gymnodinoid sequences, including *G. catenatum*, in both cores (MCS3-T2; GC2S1). Inshore, presence was detected from the most recent sediments (MCS1 2 cmbsf, ∼2 years ago, *i.e.,* 2015), in contrast to the offshore core where they were detected from 5 and 20 cmbsf (∼100 and ∼700 years ago, respectively), and sporadically throughout GC2S1 up to 209 cmbsf (∼7 638 years ago). In 1987 (30 years before the sediment core collection), *G. catenatum* was reported as representing 1% of total dinoflagellate cysts in surface sediments of Spring Bay, close to the inshore core site MCS3 (Bolch and Hallegraeff, 1990). The closest sample we analysed to this was MCS3-T2 15 cmbsf (∼30 years ago), however, no *G. catenatum sed*aDNA was detected, raising the possibility that any *G. catenatum sed*aDNA preserved was too low to be detected. In future, more thorough sequencing efforts could resolve this issue.

After refining species assignments using the new *Gymnodinium-*only reference database and detailed read verification, we detected *G. catenatum sed*aDNA sporadically through core GC2S1 (5, 20, 50, 209 cmbsf, corresponding to ∼100, ∼700, ∼2 300, and ∼7 638 years ago; deepest sample excluded). This extends the likely presence of this species in the region well beyond the proposed 1970s introduction based on analysis of resting cyst abundance in dated cores from the neighbouring Huon River (McMinn et al., 1997). Instead the *sed*aDNA approach indicates *G. catenatum* is more likely to be endemic but previously cryptic. While rDNA-ITS sequence polymorphisms indicate a link between Australasian populations and the Seto Inland Sea in Japan (Bolch and DeSalas, 2007), our findings favour proposed alternative hypotheses that recent dispersal between these areas could equally have been from Australasia to Japan (Bolch and DeSalas, 2007), or, more likely, the rDNA polymorphisms are part of a wider global biogeographical pattern of natural dispersal over many thousands of years (John et al. 2018). While the number of verified *G. catenatum* reads was very low (9 reads), their unambiguous identification highlights the known problem of overlooking low-cyst producing cryptic species via microscopy. In a companion palynological survey of the same sediment samples only 3 microreticulate cysts were found out of a total of 4 273 dinocysts counted, and was confined to 0 – 30 cmbsf of MCS3-T2 (inshore; Paine et al, in review).

### 4.4 Noctiluca scintillans

The HABbaits1 approach has revealed the presence of *N. scintillans* inshore of Maria Island during the last ∼30 years, as well as traces of this dinoflagellate in recent sediments offshore (0 – 1.5 and 5 – 6.5 cmbsf; <100 years old). Inshore, the highest relative abundances of *N. scintillans* were detected in recent sediments (from the year 2010 onwards; 6 – 7.5 and 12 – 13.5 cmbsf), indicating blooms occurred within the last 21 years. This is consistent with observational records that *N. scintillans* was first detected in Tasmania in 1994 and from 2010 onwards blooms have increased in both frequency and intensity (Hallegraeff et al., 2019).

To our knowledge, this is the first authenticated *N. scintillans sed*aDNA from a coastal ecosystem, demonstrating the sensitivity of the HABbaits1 approach for detecting plankton community change of fragile non-fossilising species from the marine sediment record. The presence of *N. scintillans sed*aDNA in recent sediments at the offshore site confirmed that its DNA has been preserved at the seafloor, after sinking through a 104 m water column. The absence of *N. scintillans sed*aDNA from older sediments (>100 years) at the offshore site (excluding the presumably contaminated deepest sample of GC2S1), indicates that DNA of this soft-bodied, non-cyst-forming dinoflagellate likely does not preserve well and may pose limitations to its analysis using *sed*aDNA records (see also Supplementary Material Note 6, Supplementary Material Fig. 6). Future investigations could target sediment cores from Sydney, where this species has been present closer to shore for longer, and the rate of damage of *N. scintillans sed*aDNA might be estimated over longer time scales. The application of similar methods to the putative range expansion of green *Noctiluca* (with green algal symbionts) into the Arabian Sea (Gómez et al., 2014) would also be of considerable interest.

## Supporting information

Supplementary Material Armbrecht et al. Maria Island dinoflagellate sedaDNA

## Acknowledgements

We are grateful to Brian Brunelle from Arbor Biosciences, USA, for his expert assistance with designing the HABbaits1 capture array. We thank Oscar Estrada-Santamaria, Steve Richards, Holly Heiniger, Nicole Moore and Steve Johnson for their help and advice during DNA extractions, library preparation, and hybridization capture. We are grateful to Raphael Eisenhofer, Vilma Pérez, Yassine Souilmi, Yichen Liu, and Ron Hübler for their help with bioinformatic analyses. We thank the Marine National Facility, the crew of RV *Investigator* voyage IN2018_T02 and the scientific voyage team for their support during field work [2018 MNF Grant H0025318]. We thank Mr Shikder Saiful Islam (IMAS, University of Tasmania) for generating additional rRNA gene sequencing from microreticulate species of *Gymnodinium*. We thank Henk Heijnis (ANSTO) for logistical support and his contributions to secure funding for this project. This study was funded through the Australian Research Council [ARC Discovery Project DP170102261]. A.C. was funded by ARC Laureate Fellowship FL140100260. LA was supported by an Australian Research Council (ARC) Discovery Early Career Researcher (DECRA) Fellowship (DE210100929).

## Author contributions

L.A. designed and carried out *sed*aDNA laboratory work, bioinformatic analyses and coordinated the manuscript writing. G.H., C.B., and C.W. collected and provided the core samples. B.P. carried out dinocysts analyses. L.A., G.H. and B.P. wrote the first draft of the manuscript. C.B. led the supplementary *Gymnodinium* rRNA gene sequencing, carried out reference sequence alignments, *sed*aDNA fragment mapping, species assignment validation and additional figures, and revised subsequent manuscript drafts. C.W. provided sediment core dating. A.C. contributed guidance on hybridization capture design and ancient DNA analyses. A.C., G.H., A.M. and C.B. developed the conceptual approach and secured funding for the project. All co-authors contributed to writing, data interpretation, comment and revision on manuscript drafts, and final editing before submission for publication.

## Abbreviations

harmful algal blooms: HAB
heuristic operations for pathogen screening: HOPS
hydrofluoric acid: HF
internally transcribed spacer: ITS
Paralytic Shellfish Poisoning: PSP
Paralytic Shellfish Toxin: PST
sedimentary ancient DNA: *seda*DNA
small subunit ribosomal rRNA: SSU
large subunit ribosomal rRNA: LSU

