## Supplementary Material Armbrecht et al. Maria Island dinoflagellate sedaDNA for "Recovering sedimentary ancient DNA of harmful dinoflagellates off Eastern Tasmania, Australia, over the last 9 000 years"

Linda Armbrecht, Bradley Paine, Christopher J.S. Bolch, Alan Cooper, Andrew McMinn, Craig Woodward and Gustaaf Hallegraeff


| **Table of content** | **Page** |
|---|---|
| Supplementary Material Note 1: Age model | 2 |
| Supplementary Material Figure 1: Age model | |
| Supplementary Material Table 1: Summary of ages | 3 |
| Supplementary Material Note 2: Shotgun rarefied | |
| Supplementary Material Figure 2: Total abundance of Dinophyceae (semi-quantitative) | 4 |
| Supplementary Material Note 3: *Alexandrium* spp., *Gymnodinium* spp. and *Noctiluca scintillans* assignment specificity testing | 5 |
| Supplementary Material Table 2: Dummy run sequences | 5 |
| Supplementary Material Figure 3: Dummy run short sequence specificity | 8 |
| Supplementary Material Note 4: Mapping of Gymnodinoid *sed*aDNA to reference sequence alignments | 9 |
| Supplementary Material Figure 4: Mapping of *Gymnodinium catenatum sed*aDNA reads to reference sequences. | 9 |
| Supplementary Material Figure 5: Sequence divergence and closest species match of gymnodinoid *sed*aDNA fragments. | 10 |
| Supplementary Material Table 3: *Gymnodinium sed*aDNA verification | 11 |
| Supplementary Material Note 5: Re-running of reads identified as *G. catenatum* against in-house *Gymnodinium* datanase | 11 |
| Supplementary Material Table 4: Re-assigned reads to *Gymnodinium* spp. | 12 |
| Supplementary Material Note 6: Sequence length analysis of cyst vs. non-cyst formers | 12 |
| Supplementary Material Figure 6: *sed*aDNA fragment lengths of dinoflagellates in Shotgun and HABbaits1 | 12 |
| References | 13 |

### Supplementary Material Note 1: Age model

The age-depth model for core MCS3-T2 and MCS2-T6 was based on eight and six lead-210 measurements, respectively. The age-depth model for core GC2S1 was based on seven lead-210 and three radiocarbon measurements. A Bayesian age-depth model for each site based on lead-210 and radiocarbon measurements was constructed using the rbacon (Blaauw et al., 2019) software in on the R platform (R Core Team, 2013) with the SHCal20 curve for radiocarbon age calibration (Hogg et al., 2020) (Supplementary Material Fig. 1A,B, Supplementary Material Table 1). It is common for surface sediment to be lost from gravity cores due to the nature of the technique, where the core is tipped horizontally after collection, which can lead to loss of unconsolidated surface sediment. We used the lead-210 activity profiles from GC2S1 and MCS1-T6 to determine the amount of material lost (Supplementary Material Fig. 1C), and estimate that 3.5 cm is missing from the top of GC2S1 which represents the last ca. 30 years. This missing time is represented in the top of MCS1-T6.

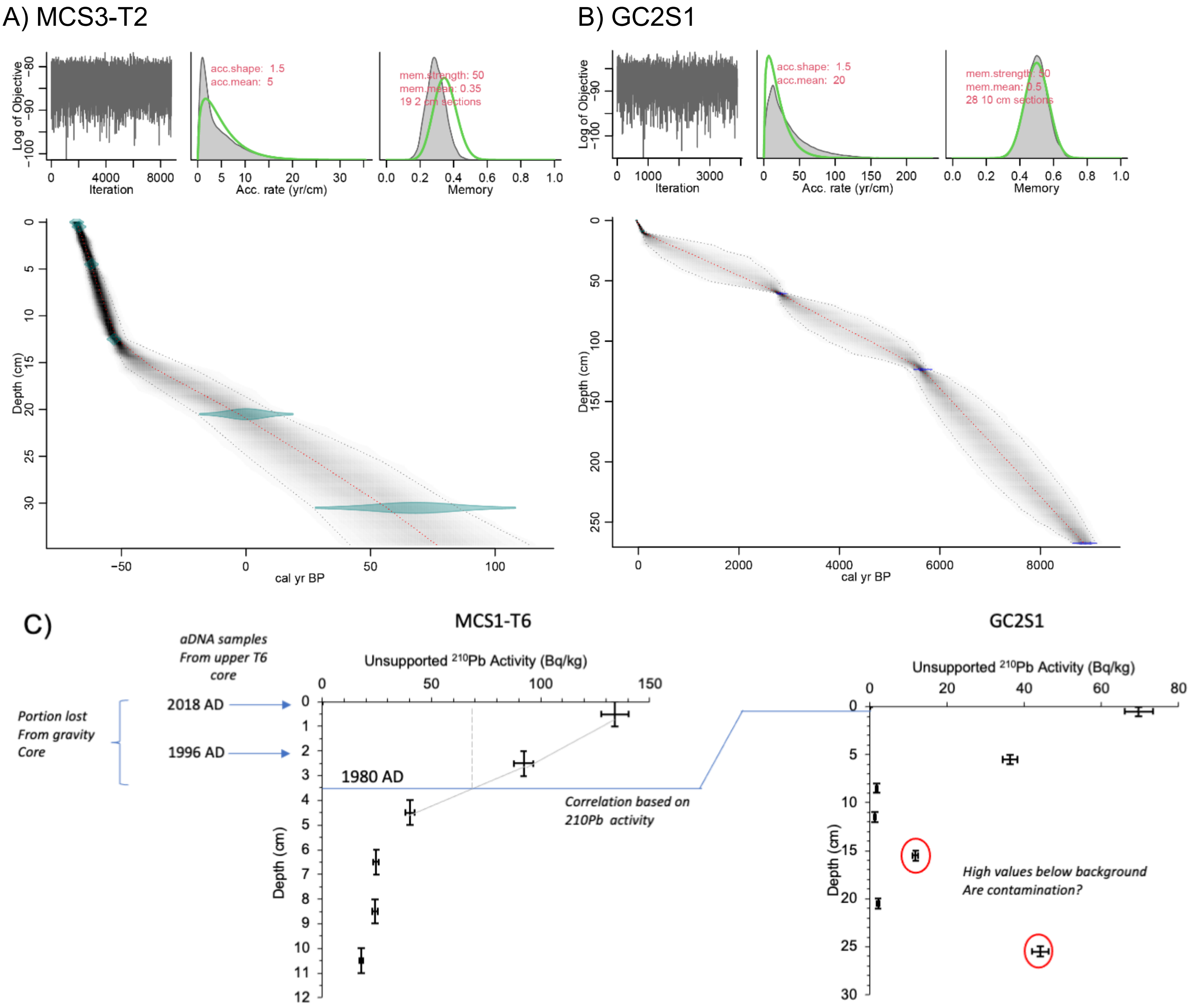


**Supplementary Material Figure 1: Age-depth models for Maria Island coring sites.** Age-depth model for A) MCS3-T2, and B) GC2S1. For ages of depths sampled for dinocysts and sedimentary ancient DNA (*seda*DNA) see Supplementary Material Table 1. C) Correlation between multicore MCS1-T6 (left) and the top of the offshore gravity core GC2S1 (right) based on lead-210 activity, revealing that 3.5 cm is missing from the top of GC2S1 (~30 years).

**Supplementary Material Table 1: Summary of ages assigned to specific sampling depths at MCS3-T2 and GC2S1.** Listed are all depths that were sampled for sedimentary ancient DNA (*seda*DNA) analyses. Ages are assigned in years relative to the year 1950 based on the model (*i.e.,* negative values indicate years after, and positive values before 1950). To facilitate interpretation in the context of arrival and spread of harmful dinoflagellate taxa off Tasmania, the age of sediments relative to the year 2017 is also provided. Note that 3.5 cm are missing in GC2S1 (~30 years).

| MCS3-T2 | | | | | | GC2S1 | | | | | |
|---|---|---|---|---|---|---|---|---|---|---|---|
| Depth (cmbsf) | Min (years BP) | Max (years BP) | Median (years BP) | Mean (years BP) | *Years ago (rel. to 2017)* | Depth (cmbsf) | Min (years BP) | Max (years BP) | Median (years BP) | Mean (years BP) | *Years ago (rel. to 2017)* |
| 0.5 | -69 | -66 | -67 | -67 | *0* | 1 | -43 | -16 | -29 | -29 | *38* |
| 2.5 | -67 | -62 | -65 | -65 | *2* | 6 | 10 | 64 | 36 | 36 | *103* |
| 4.5 | -64 | -60 | -62 | -62 | *5* | 11 | 65 | 185 | 120 | 122 | *189* |
| 6.5 | -63 | -57 | -60 | -60 | *7* | 16 | 119 | 868 | 344 | 381 | *448* |
| 12.5 | -54 | -48 | -52 | -52 | *15* | 21 | 172 | 1,504 | 575 | 642 | *709* |
| 15.5 | -48 | -22 | -39 | -38 | *29* | 26 | 332 | 1,748 | 886 | 924 | *991* |
| 20.5 | -19 | 13 | -3 | -3 | *64* | 31 | 446 | 2,211 | 1,163 | 1,203 | *1,270* |
| 25.5 | 3 | 54 | 27 | 28 | *95* | 36 | 695 | 2,330 | 1,457 | 1,464 | *1,531* |
| 30.5 | 27 | 85 | 57 | 56 | *123* | 41 | 858 | 2,535 | 1,737 | 1,726 | *1,793* |
| 34.5 | 42 | 114 | 76 | 77 | *144* | 46 | 1,174 | 2,631 | 2,035 | 1,994 | *2,061* |
| | | | | | | 51 | 1,429 | 2,786 | 2,342 | 2,264 | *2,331* |
| | | | | | | 56 | 2,142 | 2,832 | 2,580 | 2,548 | *2,615* |
| | | | | | | 61 | 2,748 | 2,962 | 2,816 | 2,824 | *2,891* |
| | | | | | | 66 | 2,810 | 3,383 | 3,009 | 3,033 | *3,100* |
| | | | | | | 71 | 2,863 | 3,890 | 3,194 | 3,245 | *3,312* |
| | | | | | | 76 | 2,976 | 4,125 | 3,442 | 3,475 | *3,542* |
| | | | | | | 81 | 3,062 | 4,533 | 3,661 | 3,705 | *3,772* |
| | | | | | | 86 | 3,247 | 4,688 | 3,914 | 3,928 | *3,995* |
| | | | | | | 91 | 3,371 | 4,965 | 4,150 | 4,153 | *4,220* |
| | | | | | | 96 | 3,621 | 5,102 | 4,415 | 4,395 | *4,462* |
| | | | | | | 101 | 3,786 | 5,339 | 4,671 | 4,635 | *4,702* |
| | | | | | | 106 | 4,071 | 5,412 | 4,909 | 4,859 | *4,926* |
| | | | | | | 111 | 4,272 | 5,562 | 5,150 | 5,082 | *5,149* |
| | | | | | | 116 | 4,861 | 5,611 | 5,338 | 5,307 | *5,374* |
| | | | | | | 121 | 5,248 | 5,721 | 5,543 | 5,524 | *5,591* |
| | | | | | | 124 | 5,355 | 5,787 | 5,624 | 5,611 | *5,678* |
| | | | | | | 125 | 5,383 | 5,827 | 5,651 | 5,640 | *5,707* |
| | | | | | | 130 | 5,495 | 6,165 | 5,767 | 5,785 | *5,852* |
| | | | | | | 135 | 5,575 | 6,299 | 5,882 | 5,898 | *5,965* |
| | | | | | | 140 | 5,627 | 6,532 | 5,984 | 6,008 | *6,075* |
| | | | | | | 145 | 5,711 | 6,668 | 6,095 | 6,120 | *6,187* |
| | | | | | | 150 | 5,759 | 6,844 | 6,209 | 6,233 | *6,300* |
| | | | | | | 155 | 5,853 | 6,961 | 6,326 | 6,345 | *6,412* |
| | | | | | | 160 | 5,928 | 7,122 | 6,438 | 6,457 | *6,524* |
| | | | | | | 165 | 6,019 | 7,225 | 6,554 | 6,569 | *6,636* |
| | | | | | | 170 | 6,091 | 7,347 | 6,668 | 6,681 | *6,748* |
| | | | | | | 175 | 6,199 | 7,449 | 6,786 | 6,796 | *6,863* |
| | | | | | | 180 | 6,274 | 7,611 | 6,905 | 6,911 | *6,978* |
| | | | | | | 185 | 6,378 | 7,693 | 7,019 | 7,021 | *7,088* |
| | | | | | | 190 | 6,460 | 7,816 | 7,132 | 7,130 | *7,197* |
| | | | | | | 195 | 6,577 | 7,910 | 7,244 | 7,241 | *7,308* |
| | | | | | | 200 | 6,671 | 8,004 | 7,358 | 7,352 | *7,419* |
| | | | | | | 205 | 6,785 | 8,092 | 7,472 | 7,462 | *7,529* |
| | | | | | | 210 | 6,868 | 8,212 | 7,587 | 7,571 | *7,638* |
| | | | | | | 215 | 7,004 | 8,290 | 7,701 | 7,681 | *7,748* |
| | | | | | | 220 | 7,103 | 8,399 | 7,809 | 7,791 | *7,858* |
| | | | | | | 225 | 7,245 | 8,477 | 7,915 | 7,905 | *7,972* |
| | | | | | | 230 | 7,356 | 8,567 | 8,031 | 8,018 | *8,085* |
| | | | | | | 235 | 7,523 | 8,634 | 8,144 | 8,129 | *8,196* |
| | | | | | | 240 | 7,614 | 8,722 | 8,259 | 8,240 | *8,307* |
| | | | | | | 245 | 7,817 | 8,779 | 8,367 | 8,351 | *8,418* |
| | | | | | | 250 | 7,941 | 8,858 | 8,480 | 8,463 | *8,530* |
| | | | | | | 255 | 8,174 | 8,912 | 8,587 | 8,579 | *8,646* |
| | | | | | | 260 | 8,329 | 8,998 | 8,704 | 8,696 | *8,763* |
| | | | | | | 265 | 8,560 | 9,060 | 8,799 | 8,804 | *8,871* |
| | | | | | | 268 | 8,651 | 9,117 | 8,878 | 8,878 | *8,945* |

**Supplementary Material Note 2: Sedimentary ancient DNA - Shotgun rarefied**

Normalising the shotgun data (i.e., rarefying/subsampling to 2.2 Mio reads per sample) reduced the number of reads assigned to Dinophyceae to a total of 148. As expected, rarefication led to the detection of less reads of harmful Dinophyceae taxa, with *Gymnodinium* only resolved to genus level in MCS3-T2 and GC2S1, and only one *Alexandrium* sequence detected in MCS3-T2 (Supplementary Material Fig. 2). In GC2S1 the read numbers showed some cyclicity throughout the core, suggesting relatively high Dinophyceae abundance (up to 10 reads per sample) between 189 and 240.5 cmbsf (~7,100 – 8,300 years ago), and 50 - 56.5 cmbsf (2,300 – 2,600 years ago) and lower read numbers in between (Supplementary Material Fig. 2). This coincided with slightly elevated *Gymnodinium* spp. read numbers at 239 cmbsf (8,300 years ago) and 50 cmbsf (~2,300 years ago), although this should be interpreted with caution due to low read numbers (Supplementary Material Fig. 2).

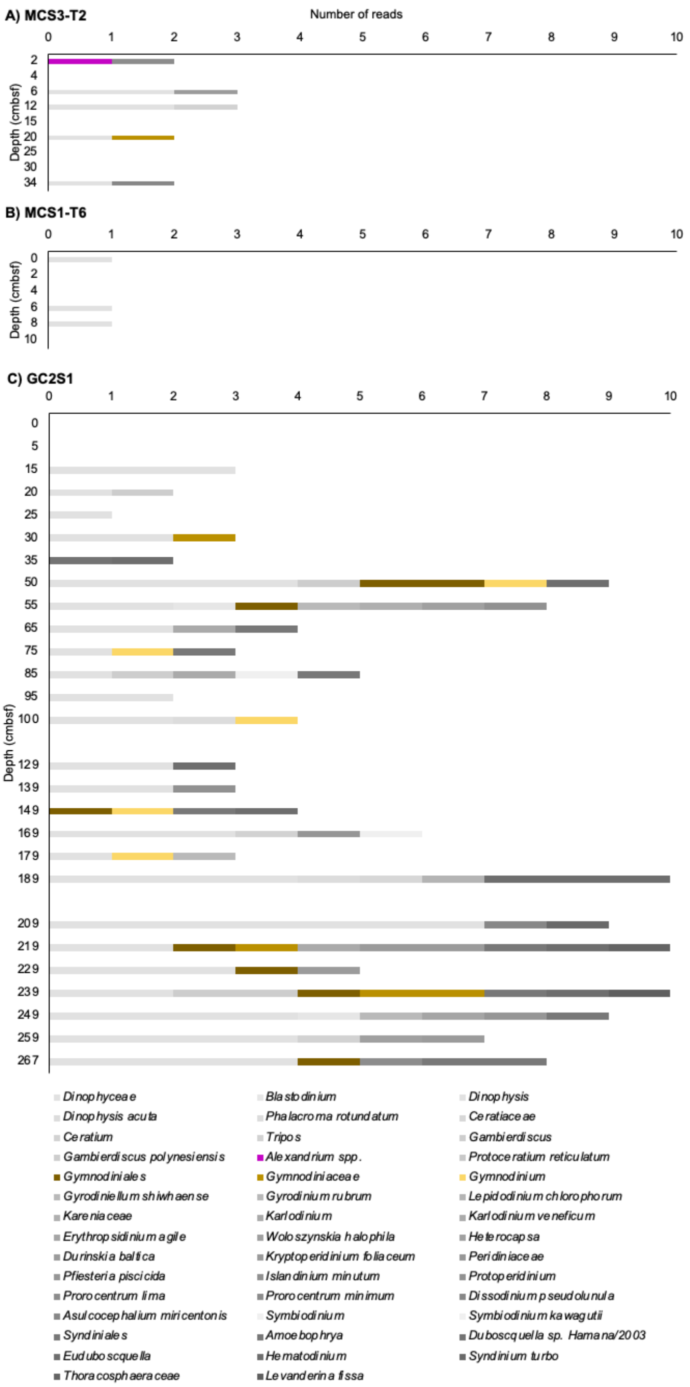


**Supplementary Material Figure 2: Total abundance of Dinophyceae (semi-quantitative).** Dinophyceae detected in rarefied Shotgun data at Maria Island coring sites A) MCS3-T2, B) MCS1-T6, and C) GC2S1.

**Supplementary Material Note 3: *Alexandrium* spp., *Gymnodinium* spp. and *Noctiluca scintillans* assignment specificity testing**

To test the reliability of assignments made to *Alexandrium* spp., *Gymnodinium* spp., and *N. scintillans,* we created three dummy samples (fasta-files) containing SSU, LSU, and ITS-5.8S-ITS and COI sequences of *G. catenatum, G. microreticulatum, G. nolleri, G. trapeziforme, G. inusitatum*, *A. tamarense* Group 1 (*A. catenella*), *A. tamarense* Group 2 (*A. mediterraneum*), *A. tamarense* Group 3 (*A. tamarense*), *A. tamarense* Group 4 (*A. pacificum*), *A. australiense*, *A. catenella*, and *N. scintillans* using sequences downloaded from NCBI (as available on 16.10.2020 for *Gymnodinium* spp., which we tested first, and as available on 23.07.2021 for *Alexandrium* spp. and *N. scintillans*). Each sequence was included with its complete length (Supplementary Material Table 3) and split into 56 bp fragments (corresponding to the average sequence length of our filtered Shotgun data). If the last fragment was <25 bp it was excluded from the dummy sample (mimicking our minimum cut-off of 25 bp during sample data processing, see main text). We converted from .fasta to .fastq for each Dummyfile separately using the reformat option in BBMap (version 36.62-intel-2017.01-Java-1.8.0_121, command: reformat.sh in=DummyFilename.fasta out1=DummyFilename_R1.fastq out2=DummyFilename_R2.fastq qfake=25 fastareadlen=129 qout=64 addcolon=t trimreaddescription=t uniquenames=t), compressed each of the resulting fastq-files (gzip) and aligned against the NCBI Nucleotide database using MALT (Herbig et al., 2016) (see main text). Additionally, we combined the fastq-files of our dummy sample with a filtered Shotgun sample that had no assigned *Gymnodinium* spp*.*, *Alexandrium* spp. or *N. scintillans* (GC2S1 5 - 6.5 cm), and re-run the alignment. The latter was performed to assess the accuracy of assignments of the dummy sequences within a complex sequence background.

**Supplementary Material Table 2:** Details of *Alexandrium* spp*.*, *Gymnodinium* spp. and *N. scintillans* sequences in dummy samples.

| Species | Accession | Gene | Length (bp) |
|---|---|---|---|
| *G. microreticulatum* | KF234058.1 | ITS1, partial sequence; 5.8S ribosomal RNA gene, complete sequence; ITS2, partial sequence | 575 |
| *G. microreticulatum* | AB265964.1 | 18S rRNA | 1,755 |
| *G. microreticulatum* | AY036078.1 | large subunit ribosomal RNA | 692 |
| *G. nolleri* | AM998534.1 | ITS1, 5.8S rRNA, ITS2 | 622 |
| *G. nolleri* | AF200673.1 | large subunit ribosomal RNA | 974 |
| *G. inusitatum* | KF234061.1 | ITS1, partial sequence; and 5.8S rRNA gene and ITS2, complete sequence | 657 |
| *G. inusitatum* | KF234072.1 | large subunit ribosomal RNA partial | 715 |
| *G. catenatum* | AF022193.1 | 18S rRNA complete | 1,804 |
| *G. catenatum* | MT659395.1 | ITS1, partial sequence; 5.8S rRNA and ITS2, complete sequence; LSU, partial sequence | 642 |
| *G. trapeziforme* | EF192414.1 | large subunit ribosomal RNA gene, partial sequence | 940 |
| *Alexandrium tamarense Group 1 (A. catenella)* | KF646464.1 | 18S ribosomal RNA gene, partial sequence; ITS1, 5.8S ribosomal RNA gene, and ITS 2, complete sequence; 28S ribosomal RNA gene, partial sequence | 1,405 |

| | | | |
|---|---|---|---|
| *Alexandrium tamarense Group 2 (A. mediterraneum)* | KF646509.1 | ITS 1, partial sequence; 5.8S ribosomal RNA gene and ITS 2, complete sequence; 28S ribosomal RNA gene, partial sequence | 1,379 |
| *Alexandrium tamarense Group 3 (A. tamarense)* | KF646463.1 | 18S ribosomal RNA gene, partial sequence; ITS 1, 5.8S ribosomal RNA gene, ITS2, complete sequence; 28S ribosomal RNA gene, partial sequence | 1,400 |
| *Alexandrium tamarense Group 4 (A. pacificum)* | KF646528.1 | SSU gene, partial sequence | 1,762 |
| *Alexandrium tamarense Group 4 (A. pacificum)* | KF646353.1 | ITS 1, partial sequence; 5.8S ribosomal RNA gene and ITS 2, complete sequence; 28S ribosomal RNA gene, partial sequence | 1,378 |
| *A. australiense* | KF908802.1 | 18S ribosomal RNA gene, partial sequence | 1,672 |
| *A. australiense* | KF985181.1 | SxtA (sxtA) gene, partial cds | 652 |
| *A. australiense* | KF908810.1 | 28S ribosomal RNA gene, partial sequence | 603 |
| *A. australiense* | KF908817.1 | ITS 1, partial sequence; 5.8S ribosomal RNA gene, complete sequence; ITS 2, partial sequence | 497 |
| *A. catenella* | KJ879220.1 | ITS1, partial sequence; 5.8S ribosomal RNA gene, complete sequence; ITS 2, partial sequence | 491 |
| *A. catenella* | KJ879232.1 | large subunit ribosomal RNA gene, partial sequence | 889 |
| *A. catenella* | KF164489.1 | SxtA4 (sxtA4) gene, partial sequence | 595 |
| *A. catenella* | GQ501122.1 | *Alexandrium catenella* voucher CS-798/03 COI gene, partial sequence; mitochondrial | 494 |
| *N. scintillans* | GQ380592.1 | 18S ribosomal RNA gene, partial sequence; ITS 1, 5.8S ribosomal RNA gene, ITS2, complete sequence; 28S ribosomal RNA gene, partial sequence | 3,576 |

*Alexandrium* species

Our results show that all assignments of our *Alexandrium* spp. dummy sample were made to *Alexandrium* on genus level (Supplementary Material Fig. 3A). This was independent of the gene used, whereby none of the *A. catenella* COI sequences (GQ501122.1) were assigned. One sequence each was misassigned to Apicomplexa and Apocrita (Supplementary Material Fig. 3A), and in both cases these were 56 bp fragments of the 18S rRNA gene, partial sequence (fragment 16 and fragment 10 of KF908802.1 *A. australiense*, respectively). These results show that *Alexandrium* sequences of the SSU, LSU, ITS1, ITS2, and SxtA could not confidently be assigned to species level, thus metagenomic sequencing data containing short reads for this genus should be interpreted on genus level.

One read each was misassigned to Apicomplexa and Apocrita, and in both cases these were short 56 bp 18S rRNA fragments (KF908802.1 *A. australiense* fragment 16, and KF908802.1 *A. australiense* fragment 10, respectively). The fragment misassigned to Apocrita was similar to a short 18S rRNA fragment of *N. scintillans* that was misassigned to the same Apocrita NCBI reference sequence (>AF436016.1|tax|177228| *Lepidopa californica* 18S ribosomal RNA gene, partial sequence, Length = 1696, see *N. scintillans* section below). This means that when Apocrita are identified alongside *Alexandrium* spp. in metagenomic data containing short reads based on 18S rRNA, the presence of Apocrita should be interpreted with caution.

*Gymnodinium* species

Not all sequences were assigned to species level but instead to higher taxonomic levels (such as *Dinophyceae, Gymnodinium*, Supplementary Material Fig. 3B). All assignments of *Gymnodinium* sequences that were achieved at species level were correct, except the 18S rRNA sequence included for *G. catenatum* (AF022193.1). Neither the complete original sequence nor any of its 56 bp fragments was assigned to *G. catenatum*, instead, the complete sequence was assigned to "Gymnodiniaceae", while the fragments were distributed across the higher taxonomic levels as well as misassigned to *Durinskia baltica, Eimeria* and Streptophytina (Supplementary Material Fig. 3B). This indicates that 18S rRNA sequences of *G. catenatum* cannot confidently be assigned to species level. Two fragments of the 18S rRNA sequence of *G. microreticulatum* were erroneously assigned to *Symbiodinium* sp. AW-2009 and Membracoidea, respectively (Supplementary Material Fig. 3B). The sequence that was misassigned to *Symbiodinium* was misassigned to the exact same NCBI reference sequence as one *N. scintillans* dummy sequence (FN552048.1|tax|675000| *Symbiodinium* sp. AW-2009 partial 18S rRNA gene, isolate 1, Length = 872). Our results suggest that assignments of Gymnodinoids to species level based on 18S rRNA sequences should be interpreted with caution, and, in particular, also the simultaneous detection of *Symbiodinium* (see *N. scintillans* below).

*Noctiluca scintillans*

Most sequences included in our *N. scintillans* dummy sample were correctly assigned to this species, and mostly correctly back-mapped to the exact NCBI sequence we used to create our dummy sample (GQ380592.1). Some reads were assigned to a different *N. scintillans* 18S rRNA reference sequence available on NCBI (>AF022200.1|tax|2966| *Noctiluca scintillans* 18S ribosomal RNA gene, complete sequence), however, the species-level identification was correct (Supplementary Material Fig. 3C).

One read each was misassigned to *Symbiodinium* and Apocrita (short 56 bp 18S rRNA fragments 9 and 11, respectively). The fragment that was misassigned to *Symbiodinium* aligned to the same NCBI reference sequence (FN552048.1|tax|675000| *Symbiodinium* sp. AW-2009 partial 18S rRNA gene, isolate 1, Length = 872) as the fragment misassigned to *Symbiodinium* in our *Gymnodinium* dummy sample (see above). The alignments were between position 477 and 422, and position 469 and 414 of the *Symbiodinium* reference sequence FN552048.1 for the *Noctiluca* and *Gymnodinium* dummy sample fragment, respectively (*i.e.,* nearly the same positions). Similarly, the fragment that was misassigned to Apocrita in our *Noctiluca* dummy sample aligned to the same NCBI Apocrita reference sequence as the fragment misassigned to Apocrita in the *Alexandrium* dummy run (NCBI reference sequence AF436016.1|tax|177228| *Lepidopa californica* 18S ribosomal RNA gene, partial sequence, Length = 1696). Again, the alignments of our short *Noctiluca* and *Alexandrium* dummy sample fragments were at very similar positions of the *Symbiodinium* sequence FN552048.1 (between position 1132 and 1077, and 1145 and 1090 for the *N. scintillans* and *Alexandrium* dummy fragment, respectively). This shows that *N. scintillans* can be accurately identified on species level using short (56 bp) sequences, however, short regions within the 18S rRNA can lead to misidentification of *Symbiodinium* and *Aprocrita,* meaning that if these latter two taxa are identified in ancient metagenomic data, the results should be interpreted with caution. The same is the case when *Symbiodinium* is identified alongside *Gymnodinium* spp., and Apocrita alongside *Alexandrium* spp.

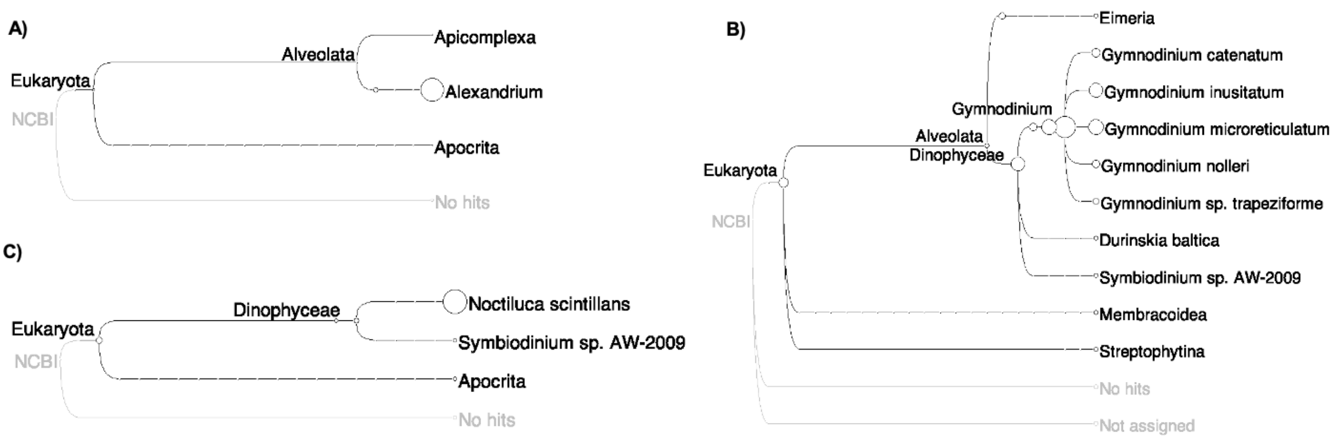


**Supplementary Material Figure 3: Dummy *Alexandrium* genus, *Gymnodinium catenatum* group. and *Noctiluca scintillans* short sequence assignment specificity.** MEGAN tree of A) *Alexandrium* spp., B) *Gymnodinium* spp. and C) *N. scintillans* identified in dummy dataset (trees created in MEGAN6CE v.18.10, Huson et al., 2016). Grey font indicates MEGAN CE's automatic function of loading a copy of the complete NCBI, displaying the taxonomy as a rooted tree.

### Supplementary Material Note 4: Mapping of gymnodinoid *seda*DNA to reference sequence alignments

Considering that species-level assignments of *Gymnodinium* spp. were generally possible based on our dummy sample analysis (above), and the fact that *Gymnodinium catenatum* was identified on species level in our data (after alignments with the NCBI nucleotide database), and its high toxicity potential, we performed additional detailed inspection of the reads assigned to *G. catenatum* in MCS3-T2 and GC2S. The *seda*DNA sequences identified as *G. catenatum* ranged from 30 to 256 bp.

Preliminary comparative alignments showed the majority of sequences mapped to the LSU-rRNA, with all but three sequences mapping to the middle section of the LSU-rRNA gene (position 4000 – 6200 of *G. catenatum* DQ785882), downstream of the D1-D2/D3 region with high taxonomic coverage. While *G. catenatum* was the closest match for most of the LSU *seda*DNA, only two other publicly available gymnodinoid sequences cover the D3-D10 region, *G. impudicum* (DQ779993), and *G. aureolum* (DQ779991). Comparative analysis of DQ779991 indicated that this sequence comes from the more distantly related genus *Karenia* therefore we excluded it from further comparisons.

The low taxon coverage in the D3-D1 region means there is high potential for reference bias. To both investigate and reduce potential bias, we generated additional sequences of the D3 to D10 region of the LSU-rRNA from two of five species *Gymnodinium* "microreticulate-group" (the group to which *G. catenatum* belongs) available to us in culture: *G. nolleri*, the most closely related species, and *G. microreticulatum* the most divergent of the group and known from the area where this study's sediment cores were collected. Using DNA extracted from cultures *G. nolleri* (strain K-0626) and *G. microreticulatum* (strain CAWD191), we amplified several overlapping fragments of the D3-D10 region of the LSU rRNA ranging from ~1,000 - 2,000 bp. PCR was carried out using combinations of forward primers D1R (Scholin et al., 1994) and D3A (Nunn et al., 1996) with reverse primers 1483R (Daugbjerg et al., 2000) GLD677 (Nishimura et al., 2013) and R8 (Chinain et al., 1999), to ensure 2x to 3x high quality sequence coverage over the D3 to D10 region. PCR was carried out using MyTaqHS polymerase and buffer system according to manufacturer's guidelines (Bioline, Australia) and thermal cycling as follows: 2 min at 95 °C, followed by 35 cycles of: 10 sec at 95 °C, 20 sec at 55 °C, and 20 sec at 72 °C. Sanger dye-terminator sequencing in both F and R directions was carried out using the amplification primers and manufacturers protocols, and sequenced on an ABI9600 sequencer at the Ramaciotti Centre for Functional Genomics (RCFG, University of NSW, Sydney). Resulting sequences were trimmed of low-quality reads, assembled using Geneious Prime 2021 and a D3-D10 region consensus established for each species using highest quality call to resolve base conflicts.

We then undertook further validation of the 352 *seda*DNA reads assigned to *Gymnodinium* from core GC2S1 (see Main Text) by mapping all gymnodinoid *seda*DNA sequences, new *G. nolleri*, *G. microreticulatum* D3-D10 sequences, and existing near full LSU-rRNA gene sequences of *G. impudicum* (DQ779993), to *G. catenatum* (DQ785882) as a reference sequence. Reference alignment and mapping used alignment/assembly algorithms implemented in the package Geneious Prime 2021 (default overlaps and word length, gaps allowed; Supplementary Material Fig. 4). The length of mapped *seda*DNA ranged from 30 bp to 256 bp (median length = 60 bp). As anticipated from the HABbaits1 design (Armbrecht et al., 2021), the majority (99%) mapped to the rRNA genes; the LSU-rRNA gene (81%), the rDNA-ITS region (14%) and the SSU-rRNA gene (5%) (Supplementary Material Table 3). Three sequences were assigned to cytochrome oxidase (CO1), a target also included in HABbaits1. While correctly assigned to other gymnodinoid species/genera, we excluded these sequences from further comparison.

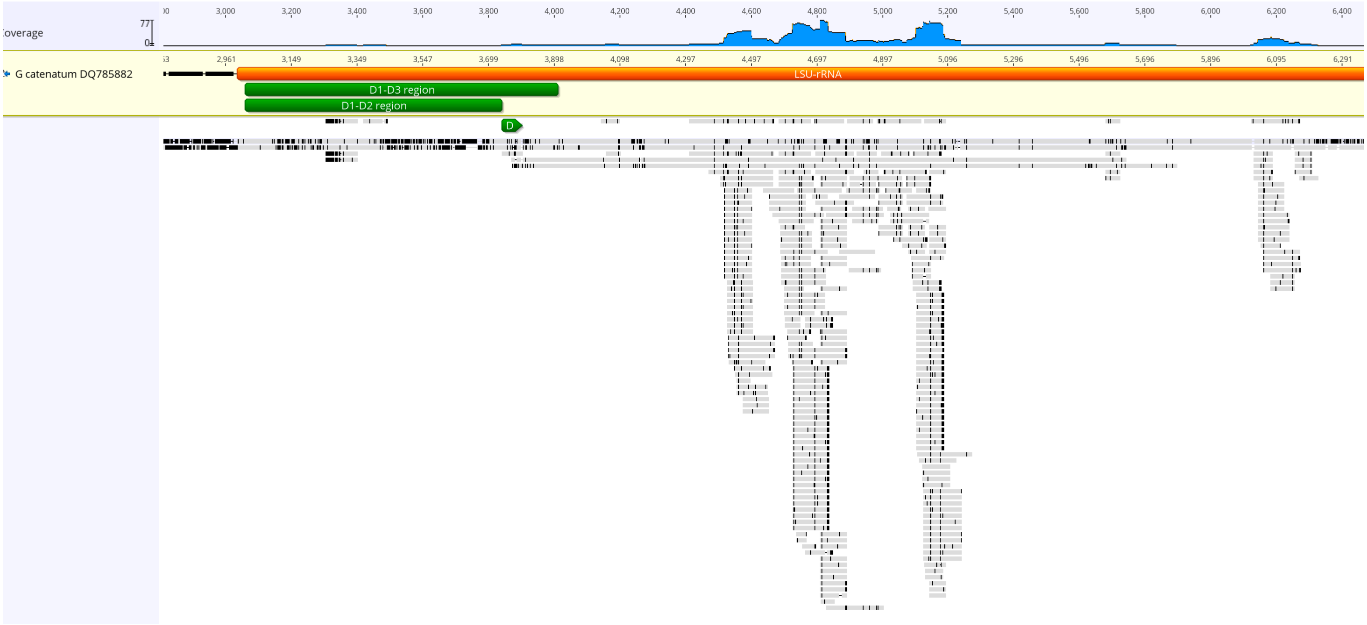


**Supplementary Material Figure 4: Mapping of *Gymnodinium catenatum seda*DNA reads to reference sequences.** Example of fragment mapping (middle section of LSU-rRNA shown) of *seda*DNA fragments to *G. catenatum* DQ785882 (reference sequence), related *Gymnodinium* sequences, and extended LSU-rRNA sequences of *Gymnodinium microreticulatum* and *G. nolleri* (Reference alignment, default parameters; Geneious Prime 2021).

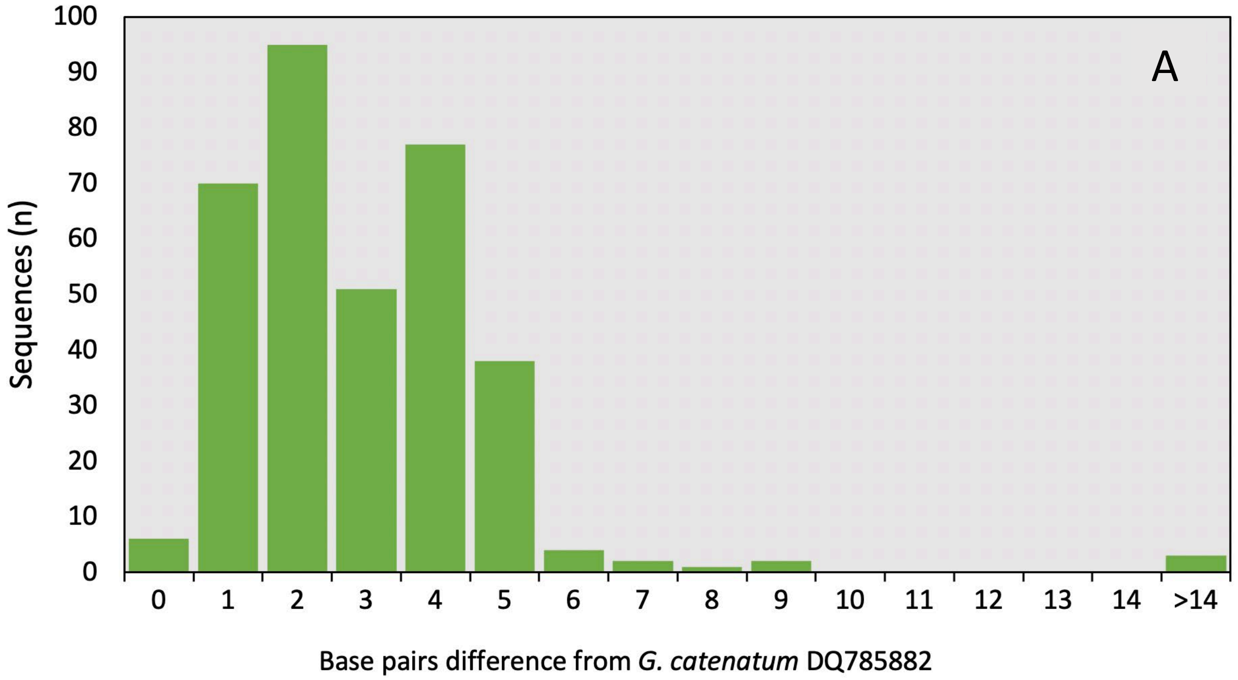


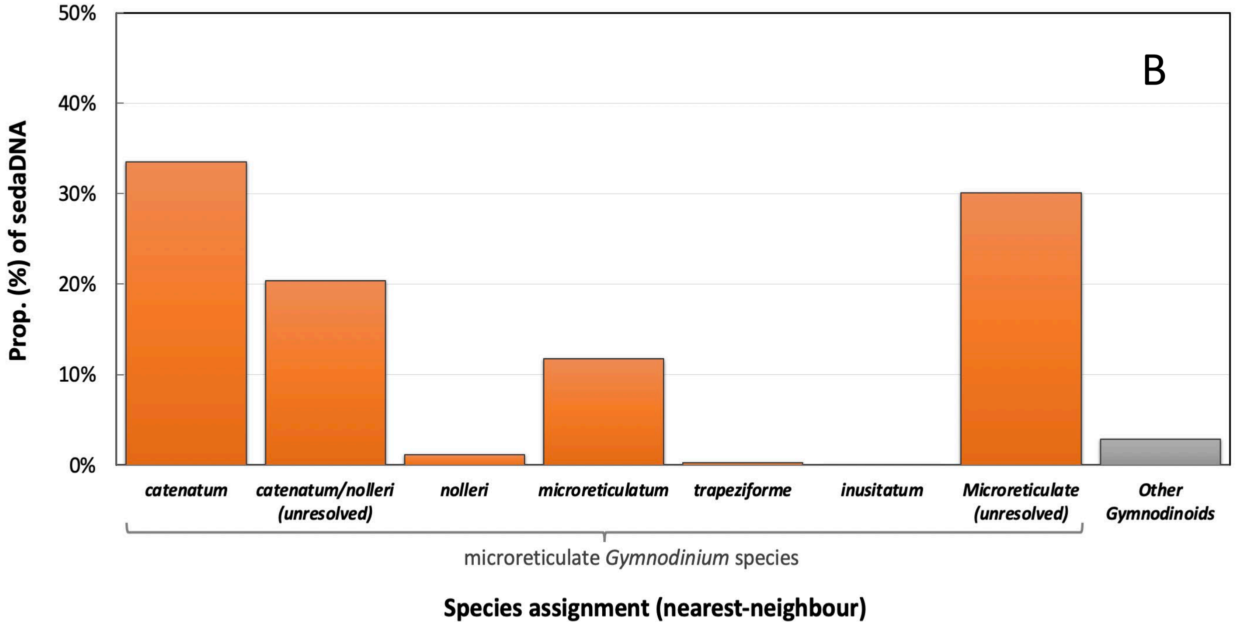


**Supplementary Material Figure 5: Sequence divergence and closest species match of gymnodinoid *sed*aDNA fragments.** A) Distribution of sequence divergence (bp) of *sed*aDNA from *G. catenatum* DQ785882. B). Proportion of fragments assigned to microreticulate-group species, microreticulate group-unresolved, and *Gymnodinium* species-unresolved (reference alignment, default parameters; Geneious Prime 2021).

**Supplementary Material Table 3: *Gymnodinium sed*aDNA verification**. *Gymnodinium* species assignments are based on closest sequence match between *sed*aDNA fragments after alignment to reference rRNA gene sequences of five known microreticulate-group species and *G. impudicum.* Also provided are the number of base mis-matches of *Gymnodinium sed*aDNA to *G. catenatum* reference sequence DQ785882.

| **Species assignment** | **Proportion of *sed*aDNA (%) and read location** | | | |
|---|---|---|---|---|
| | **LSU-rRNA (n=284)** | **SSU-rRNA (n=18)** | **5.8S rRNA (n=47)** | **Total (n=349)** |
| **Microreticulate species group** | 96.5 | 100 | 100 | 97.1 |
| *G. catenatum* | 38.0 | 44.4 | 2.1 | 33.5 |
| *G. catenatum* or *nolleri* (unresolved) | 7.4 | 38.9 | 91.5 | 20.3 |
| *G. nolleri* | 1.1 | 5.6 | | 1.1 |
| *G. microreticulatum* | 14.1 | | 2.1 | 11.7 |
| *G. trapeziforme* | | 5.6 | | 0.3 |
| *G. inusitatum* | | | | |
| Unresolved microreticulate spp. | 35.9 | 5.6 | 4.3 | 30.1 |
| **Other *Gymnodinium* species** | 3.5 | | | 2.9 |
| **Base mis-match to *G. catenatum*** | **Proportion of fragments (%)** | | | |
| | **LSU-rRNA (n=274)** | **SSU-rRNA (n=18)** | **5.8S rRNA (n=47)** | **Total (n=339)** |
| 0 | 1.8 | 0 | 2.1 | 1.8 |
| 1 | 9.5 | 22.2 | 85.1 | 21.3 |
| 2 | 28.8 | 55.6 | 12.8 | 29.0 |
| 3 | 17.5 | 16.7 | 0 | 15.5 |
| 4+ | 42.3 | 5.6 | 0 | 35.7 |

Of the reads assigned to rRNA genes, 68% were within 3 bp of exact match for *G. catenatum* reference sequence (Supplementary Material Fig. 5A; Table 3) and more than 97% were unambiguously assigned to the microreticulate-group of *Gymnodinium* species that includes *G. catenatum*. More than half (54%) of reads were closest, or equal closest match to *G. catenatum* reference sequence, with 33.5% unambiguously assigned to *G. catenatum*, and 12% to *G. microreticulatum*. Approx 20% mapped to rRNA regions with insufficient sequence variation to clearly resolve *G. catenatum* from closest relative *G. nolleri*; and 30% to regions unable to distinguish among the microreticulate-group species (Supplementary Material Fig. 5B).

**Supplementary Material Note 5: Re-running of reads identified as *G. catenatum* against an in-house *Gymnodinium* database**

We created a database of *Gymnodinium* sequences by compiling the same ten publicly available (ncbi) *Gymnodinium* sequences used in the dummy-run (Supplementary Material Table 2) and the two newly acquired *G. nolleri* and *G. microreticulatum* sequences into one .fasta-file. Next, we exported all reads (total of 352) from MEGAN CE that had been identified as *G. catenatum* after running our HABbaits1 data against the NCBI nucleotide database (Main text). We converted from .fasta to .fastq for each file separately using the reformat option in BBMap (command: reformat.sh in=Filename.fasta out=Filename.fastq), compressed each of the resulting fastq-files (gzip) and aligned against the new in-house *Gymnodinium* database using MALT (Herbig et al., 2016) with the same parameters as the NCBI and dummy runs. This resulted in 216 assigned reads, mostly identified as *Gymnodinium* spp. and to a lesser degree *G. nolleri, G. microreticulatum* and *G. catenatum* (Supplementary Material Table 4).

**Supplementary Material Table 4: Re-assigned reads to *Gymnodinium* spp.**

| Site | GC2S1 | | | | | | | | | | | MCS3-T2 | | | | | | |
|---|---|---|---|---|---|---|---|---|---|---|---|---|---|---|---|---|---|---|
| **Depth (cmbsf)** | **5** | **20** | **30** | **50** | **65** | **75** | **85** | **169** | **209** | **219** | **267** | **2** | **6** | **12** | **20** | **25** | ***Total*** | ***Proportion (%)*** |
| *Gymnodinium* | 0 | 2 | 5 | 2 | 7 | 4 | 6 | 1 | 74 | 11 | 6 | 40 | 1 | 4 | 2 | 0 | *165* | *76* |
| *G. catenatum* | 1 | 1 | 0 | 1 | 0 | 0 | 0 | 0 | 3 | 0 | 3 | 0 | 0 | 0 | 0 | 0 | *9* | *4* |
| *G. microreticulatum* | 0 | 1 | 0 | 0 | 0 | 0 | 0 | 0 | 3 | 0 | 0 | 10 | 0 | 0 | 0 | 0 | *14* | *6* |
| *G. nolleri* | 0 | 2 | 0 | 0 | 3 | 0 | 0 | 1 | 4 | 2 | 2 | 14 | 0 | 0 | 0 | 0 | *28* | *13* |

**Supplementary Material Note 6: Sequence length analysis of cyst vs. non-cyst formers**

After alignments with the NCBI Nucleotide database (see Main Text), read-length data was exported from MEGAN CE (Huson et al., 2016) for Dinophyceae and the genera *Gymnodinium*, *Alexandrium* and *Noctiluca* (separately for each coring site) to assess whether cyst-formers preserve better than non-cyst formers (assumedly reflected in longer vs. shorter read lengths, respectively). However, due to very low read numbers recovered by the shotgun data and high standard deviations of average fragment lengths per taxon, these data are provided here within the Supplementary Material.

Slight differences in *Alexandrium*, *Gymnodinium* and *Noctiluca sed*aDNA read lengths were found when compared to overall Dinophyceae read lengths in the shotgun and HABbaits1 data (Supplementary Material Fig. 7). *Gymnodinium* sequence length seemed to be slightly longer in comparison to the overall Dinophyceae sequences, which is consistent with this species being a good cyst-former, and which may benefit its *sed*aDNA preservation over time. However, due to very low number of reads recovered in the shotgun data, and high standard deviations of average fragment lengths per taxon in both shotgun and HAbbaits1 datasets, we do not consider this read-length data statistically robust. It is provided here for completeness.

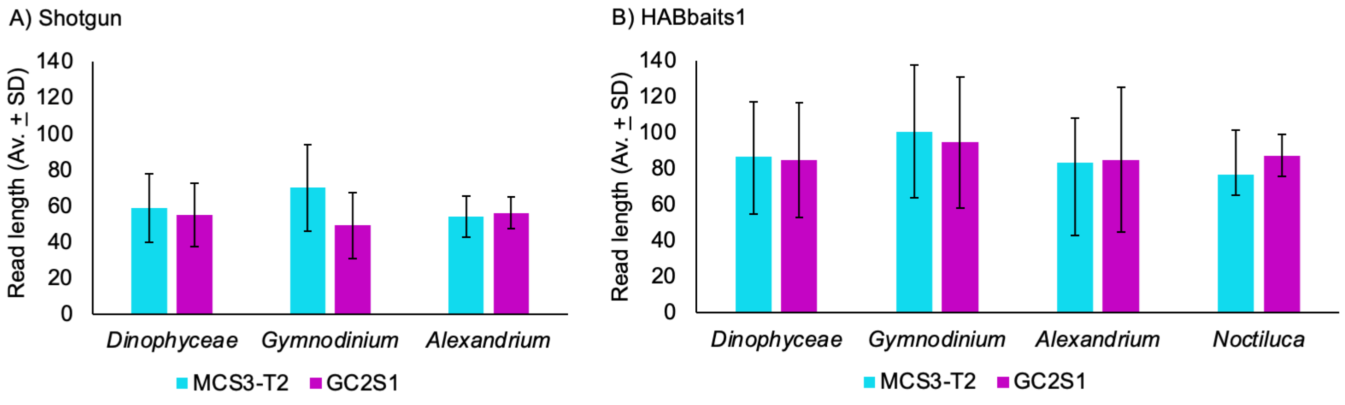


**Supplementary Material Figure 6: *sed*aDNA fragment lengths of dinoflagellates in Shotgun and HABbaits1.** Shown are the average read length (Av.) and standard deviation (SD) of sequences assigned to Dinophyceae, *Gymnodinium* (good cyst-former), *Alexandrium* (weak cyst-former) and *Noctiluca* (non-cyst-former) in A) Shotgun and B) HABbaits1. Note that we did not detect *Noctiluca* in the shotgun dataset.
